# Multi-omics integration reveals candidate genes and molecular mechanisms associated with resistance to sugarcane smut

**DOI:** 10.64898/2026.11.05.764367

**Authors:** Débora Thomazim, Ricardo José Gonzaga Pimenta, Lucélia de Fátima Santos, Aline da Costa Lima Moraes, Danilo Augusto Sforça, Luciana Oliveira Souza Anjos, Ivan Antônio dos Anjos, Felipe Brussolo da Silva, Luciana Rossini Pinto, Anete Pereira de Souza

**Author notes:** Corresponding author: Anete Pereira de Souza, Ph.D.

## Abstract

Sugarcane smut, caused by the fungus *Sporisorium scitamineum* (Syd.), is one of the main diseases affecting sugarcane and is capable of causing severe yield losses in susceptible varieties. Given the limitations of classical breeding, mainly due to the genetic complexity of sugarcane cultivars, molecular and omics approaches have gained increasing prominence for studying the genetic basis of traits such as disease resistance. In this context, we investigated sugarcane resistance to smut through the integration of association mapping methods and transcriptomics. A panel of 155 genotypes was genotyped with 33,335 SNP markers and inoculated with *S. scitamineum* generating three phenotypic variables. These data were jointly analyzed through genome-wide association analyses to identify marker-trait associations. In addition, transcriptomic analyses were performed using publicly available datasets from four studies evaluating sugarcane responses to smut. The integration of these multi-omics data enabled the identification of candidate genes involved in structural reinforcement, immune signaling, redox homeostasis, genome maintenance, and cellular adaptation. These findings provide valuable molecular resources for functional validation and breeding strategies aimed at developing sugarcane cultivars with enhanced smut resistance.

## 1. Introduction

Sugarcane (*Saccharum* spp.) is a tropical plant with high photosynthetic efficiency, belonging to the C4 group and to the Poaceae family. Due to its efficiency in synthesizing and accumulating sucrose, it currently ranks as the crop with the highest gross production worldwide [1,2]. Modern sugarcane cultivars, derived from domestication processes and plant breeding strategies, consist predominantly of interspecific hybrids between *Saccharum officinarum* and *Saccharum spontaneum*. These cultivars exhibit particularly challenging genomic characteristics, such as a high degree of polyploidy, frequent aneuploidies, and a variable number of chromosomes, making their genomes more complex than those of most crops [1,3,4]. Given these challenges, conventional breeding techniques are limited, and molecular and omics-based approaches can provide effective means to address these difficulties and contribute to a better understanding of its genome and the genetic basis of traits of interest, such as disease resistance [1].

One of the most important and prominent diseases affecting sugarcane is smut, caused by the fungus *Sporisorium scitamineum* (Syd.). The disease is difficult to prevent and causes significant economic losses, as susceptible varieties may experience losses in juice quality and yield reductions that can reach up to 100% [5,6,7]. Colonization by *S. scitamineum* results in the formation of a long black whip, which establishes itself in the primary shoots of infected plants. The extent of infection, and consequently the damage caused, depends on several factors, including the host plant degree of susceptibility, soil conditions, temperature, pathogen virulence, and teliospore concentration [7,8,9].

The primary method for smut control is the development and planting of resistant varieties [6]*. S. spontaneum* has long been recognized as an important source of disease resistance in sugarcane breeding; however, resistance to smut is not uniformly distributed among its accessions. Considerable variation in resistance levels has been reported, highlighting the importance of characterizing and selecting resistant genotypes for breeding purposes. This variability represents a challenge for breeding programs aiming to develop high-biomass cultivars, which rely on the genetic contribution of *S. spontaneum* [1,10,11]. Therefore, understanding the genetic basis of smut resistance has become essential for the efficient identification and introgression of resistance-associated alleles into elite breeding populations. Molecular tools and approaches have consequently played a key role in elucidating the mechanisms underlying resistance and accelerating cultivar development.

A few mechanisms conferring resistance to *S. scitamineum* have been identified in sugarcane through molecular approaches. A key mechanism involves a hypersensitive response in infected tissues, regulated by reactive oxygen species (ROS) and nitric oxide (NO), which leads to programmed cell death and reduced fungal colonization in resistant genotypes. This response includes an oxidative burst with direct toxic effects on the pathogen, membrane damage, cell wall modifications, antimicrobial compound synthesis, and modulation of gene expression [6,12,13,14]. Although only a limited number of resistance genes have been identified to date, genetic engineering studies have demonstrated the successful introduction of resistance-conferring genes into sugarcane, such as *thaumatin* (*PR5*), which encodes an antifungal protein, and *broad-spectrum resistance 1* (*BSR1*), a cytoplasmic kinase capable of enhancing host resistance [15,16]. Nevertheless, the molecular basis of smut resistance remains only partially understood, and the genetic factors underlying durable resistance in diverse sugarcane germplasm are still largely unexplored. Further research is therefore needed to identify additional resistance-associated loci, clarify resistance mechanisms, and support the development of more effective breeding and biotechnological strategies for smut control.

Transcriptomic approaches have also been successfully applied to the study of sugarcane smut, enabling the identification of differentially expressed genes (DEGs) between infected and control plants and advancing the understanding of host-pathogen interactions. In particular, differential expression has been observed in genes associated with phytohormone signal transduction pathways, as well as in protein metabolism and cell wall formation [17,18]. In addition, smut-infected plants have been shown to reprogram their metabolism toward energy production and synthesis of signaling molecules [19], with most DEGs associated with metabolic pathways [20]. These approaches have also enabled the comparison of *S. scitamineum* isolates with different virulence levels, revealing stronger transcriptional reprogramming in response to highly virulent strains, including modulation of transcription factors and defense-related genes [21]. Furthermore, co-expression network analyses (WGCNA) have identified resistance-associated gene modules and hub genes potentially involved in immune response and ROS metabolism [22].

Despite the advances achieved, the mechanisms conferring disease resistance in plants are complex and involve interactions among multiple alleles and genes. The repeated use of a limited number of major resistance genes can impose strong selection pressure on pathogen populations, accelerating resistance breakdown. Continuous identification and incorporation of additional resistance sources are therefore essential to strengthen durability and ensure long-term effectiveness in breeding programs [23,24]. For sugarcane, expanding the identification of molecular markers and resistance-associated genes linked to smut response is particularly important, given the continuous emergence of new virulent races capable of overcoming host defenses. In this context, the discovery of novel resistance sources remains imperative, and multi-omic approaches represent a powerful strategy to provide deeper insights into pathogen biology, host resistance mechanisms, and the development of sustainable management strategies [9,25].

In this study, we investigated sugarcane resistance to smut through the integration of association mapping and transcriptomic analyses. This approach enabled the identification of associated molecular markers with potential application in the early selection of resistant genotypes, as well as candidate genes and biological processes underlying resistance and susceptibility responses to *S. scitamineum*. By integrating genomic and transcriptional evidence, this study provided a robust multi-omic approach that advances the current understanding of sugarcane-smut interactions at the molecular level and offers valuable resources for future functional studies and resistance-oriented breeding strategies.

## 2. Material and methods

### 2.1. Plant material and genotyping

The mapping study was conducted using a panel of 155 sugarcane genotypes from the Instituto Agronômico de Campinas (IAC) (Table S1). Genotyping of 97 accessions using genotyping-by-sequencing (GBS) from this panel had been previously performed, and analyses demonstrated the existence of genetic and phenotypic variability with respect to resistance to other diseases, such as yellow leaf and mosaic, as well as to other agronomically relevant traits [26,27,28].

For the 58 genotypes that had not yet been genotyped, DNA extraction followed by GBS was performed. Total genomic DNA was extracted from young leaf tissue using the GenElute Plant Genomic DNA Miniprep Kit (Sigma-Aldrich, St. Louis, USA), with DNA integrity evaluated by electrophoresis on a 1% (w/v) agarose gel stained with ethidium bromide. The DNA concentration of each genotype was subsequently measured using a Qubit 3.0 fluorometer (Thermo Scientific, Wilmington, USA).

GBS library construction was performed based on the protocol described by Poland [29], using the restriction enzymes *PstI* and *MseI*, following the same methodology previously applied to part of the panel [26]. The library was validated using a TapeStation 4200 system (Agilent Technologies, Santa Clara, USA) and quantified by quantitative PCR (qPCR) on a CFX384 detection system (Bio-Rad Laboratories, Hercules, USA). Sequencing was carried out on a NovaSeq 6000 platform (Illumina, San Diego, USA) using an SP flow cell, in single-end mode, with a read length of 100 bp. Sequencing quality was assessed and visualized using FastQC software [30].

The TASSEL4-Poly pipeline, developed by Pereira [31] based on modifications to TASSEL software version 4.0 [32], was used for single nucleotide polymorphism (SNP) discovery in the population under study. Tags were mapped against reference sequences using Bowtie2 version 2.2.5 [33]. Three sugarcane genomes were tested as references for tag alignment: a methyl-filtered genome of the hybrid cultivar SP70-1143 [34], which has shown favorable results for SNP discovery by GBS in sugarcane [26,35,36]; and the chromosome-level genomes of two hybrid sugarcane cultivars recently made available [37,38].

Genotypic class definition was performed based on the proportion of read depth of alleles at each locus, following an approach previously employed in other studies [26,27]. Markers were first filtered using VCFtools version 0.1.13 [39], retaining biallelic polymorphisms with sequencing depth greater than 50 reads, a minor allele frequency of at least 5%, and up to 25% missing data. After this step, a single reference genome was selected for subsequent analyses, and genotypes for each marker were called assuming fixed ploidy levels of 6, 8, 10, 12, and 14, according to the proportion of allelic read depths at each locus, using the Updog package [40] in the R software environment [41]. The ploidy level yielding the highest number of high-quality markers was subsequently selected for downstream analyses.

### 2.2. Phenotyping

The assay to evaluate resistance to sugarcane smut was conducted previously [42] between March 2022 and May 2023 under both greenhouse and field conditions at the Advanced Division of Sugarcane Research and Development, from Agronomic Institute Campinas (IAC), located in Ribeirão Preto, São Paulo, Brazil.

#### 2.2.1. Greenhouse trial

A randomized complete block design with three replicates was adopted, in which each experimental plot consisted of 50 buds, totaling 150 buds per genotype. The genotypes SP80-3280 and IAC6-66 were used as resistant and susceptible controls, respectively. Twelve-month-old stalks were used to obtain individualized buds following the pre-sprouted seedling methodology [43].

Prior to planting, buds were subjected to artificial inoculation using the puncture method. The inoculum consisted of a paste prepared by homogenizing 0.20 g of viable *S. scitamineum* teliospores in a water-Tween 20 solution until a cream-like consistency was obtained. Each bud was gently punctured on both sides using a needle, preserving the germinative pole. After inoculation, the buds were placed in trays with 50 cells containing Carolina Soil Class XVI substrate and incubated in a germination chamber with relative humidity above 85% for 24 h at 28°C to allow teliospore germination, followed by an additional 24 h at 31°C to promote infection. After this period, the temperature was maintained at 35°C for approximately 5 days to accelerate bud sprouting. Subsequently, the trays were transferred to the greenhouse under sprinkler irrigation, where they remained until phenotypic evaluations.

Evaluations began two months after bud planting and the appearance of the first whips and were performed weekly up to 198 days after the beginning of the evaluations. Disease incidence, represented by the percentage of plants producing whips, was calculated by dividing the number of plants with whips by the total number of plants in each plot, according to the equation:

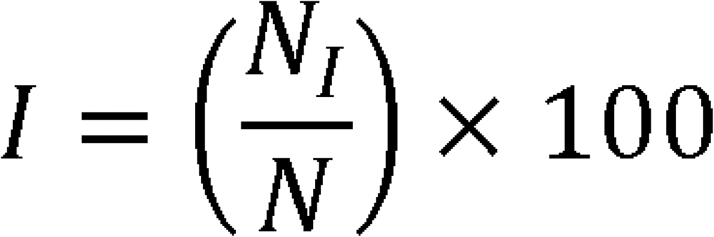

Where *I* is disease incidence, *N_I_* is the number of infected plants, and *N* is the total number of sprouted buds. To account for disease development over time, the Area Under the Disease Progress Curve (AUDPC) was also calculated according to the equation proposed by Campbell and Madden [44]:

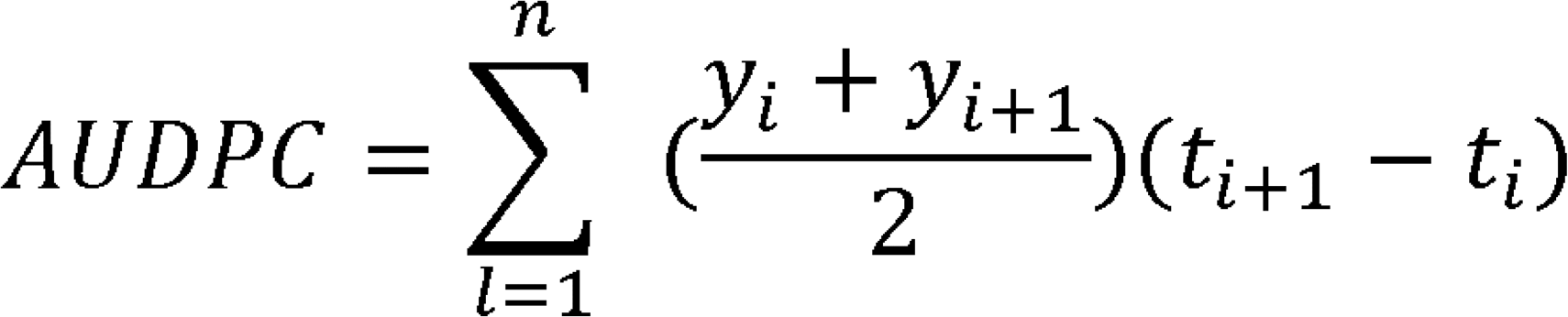

Where *n* is the total number of observations, *y* is the incidence at the *i*^th^ observation, and *t* is the time at the *i*^th^ observation. AUDPC was calculated at each evaluation until the end of the experimental assessment cycle, in May 2023.

#### 2.2.2. Field trial

The genotypes were evaluated under field conditions following the methodology described in Section 2.2.1, with the adaptations described below. The experimental design was a randomized block design with three replicates, with each plot consisting of four plants spaced 0.5 m apart within rows and 1.5 m between rows, totaling 12 plants per genotype. Prior to transplanting, plants underwent an acclimatization process for two months. Water supply relied exclusively on rainfall during the experimental period. After transplanting, disease symptoms were evaluated following the same criteria described in Section 2.2.1, during the period from November 2022 to May 2023.

### 2.3. Analysis of phenotypic data

Phenotypic data for the three evaluated traits, (i) percentage of whips in the greenhouse assay (GH), (ii) percentage of whips in the field assay, and (iii) AUDPC, were statistically analyzed using R software to assess data distribution, identify the presence of significant differences among genotypes and experimental blocks, and determine the most appropriate statistical treatment of the data.

Data normality was tested using the Shapiro-Wilk test, adopting a significance level of 5% (*p*-value < 0.05). Differences among genotypes and blocks were evaluated using the Kruskal-Wallis test, also at a 5% significance level. In addition, relationships among the three traits were explored through Pearson correlation analyses to quantify linear association between the evaluated parameters.

### 2.4. Association analyses

Association analyses were performed using mixed linear modeling approaches implemented in the GWASpoly R package [45], following a methodology previously applied to sugarcane in other studies [26,27]. According to a proposal by the package developers, several population structure and kinship matrices were constructed and included as covariates to determine the combination that minimized spurious associations.

Two methodologies were used to obtain population structure (Q) matrices for the panel: (i) discriminant analysis of principal components (DAPC), performed using the adegenet package [46] in R; and (ii) principal component analysis (PCA) followed by K-means clustering, conducted in R, using the elbow and silhouette methods implemented in the factoextra package [47] to determine the optimal number of clusters.

Similarly, three different kinship (K) matrices were computed for the panel: (i) a realized relationship matrix (MM^T^) [48], implemented directly in GWASpoly; (ii) a full autopolyploid matrix; and (iii) a pseudodiploid matrix, both based on Slater [49] and computed using the AGHmatrix package [50] in R.

For each combination of population structure and kinship matrices, six marker effect models available in GWASpoly were tested: general, additive, simplex dominant reference, simplex dominant alternative, diploidized general, and diploidized additive. For each model, a quantile-quantile (Q-Q) plot of marker ™log_10_(*p*-values) was generated, from which the inflation factor (λ), corresponding to the regression coefficient of the Q-Q plot, was calculated. The mean λ across all models using the same Q-K combination was calculated and used to select the best set of matrices. Once this combination was chosen, models showing appropriate *p*-value inflation (as verified by Q-Q plots) were selected. To determine the significance of associations, the false discovery rate (FDR) correction [51] was used with α = 0.05.

Based on the annotated scaffolds of the methyl-filtered genome, a 100-nt window up- and downstream of each resistance-associated SNP was extracted, generating 201-nt sequences centered on each SNP. These sequences were aligned against the genome of the R570 cultivar using BLASTn [52]. Genes present in these regions were verified using the JBrowse genome browser available in Phytozome [53], according to the R570 gene annotation. In addition to SNP annotation, this procedure enabled the identification of homologous genomic regions in the R570 genome and the annotated genes located within these regions, allowing the integration of genomic and transcriptomic information.

### 2.5. Transcriptome data

RNA-Seq data were obtained from publicly available datasets derived from four previously published studies [17,19,21,22]. These studies sampled different tissues (shoots, whips, and meristematic tissue) from distinct sugarcane genotypes under both control conditions and upon infection by *S. scitamineum* for different time periods, comprising 27 distinct treatment conditions, each supported by three biological replicates, totaling 81 samples (Table 1).

**Table 1.** Summary of the transcriptomic datasets used in the analyses. For each study, the sugarcane genotype and smut resistance phenotype, treatment and fungal isolate are presented.

| **Study** | **Genotype** | **Phenotype** | **Treatment** | **Fungal isolate** |
|---|---|---|---|---|
| Schaker *et al*. (2016) | RB925345 | Intermediately resistant | 5 DAI (Control) | - |
|  |  |  | 200 DAI (Control) | - |
|  |  |  | 5 DAI | SSC39 |
|  |  |  | 200 DAI | SSC39 |
| McNeil *et al*. (2018) | CP74-2005 | Susceptible | 48 HAI (Control) | - |
|  |  |  | 48 HAI | NA |
| Agisha *et al*. (2022) | Co 97009 | Susceptible | 2 DAI (Control) | - |
|  |  |  | 5 DAI (Control) | - |
|  |  |  | 60 DAI (Control) | - |
|  |  |  | 2 DAI | Ss97009 |
|  |  |  | 5 DAI | Ss97009 |
|  |  |  | 60 DAI | Ss97009 |
|  |  |  | 2 DAI | SsV89101 |
|  |  |  | 5 DAI | SsV89101 |
|  |  |  | 60 DAI | SsV89101 |
| Wu *et al.* (2022) | YT93-159 | Resistant | 0 DAI (Control) | - |
|  |  |  | 1 DAI | NA |
|  |  |  | 2 DAI | NA |
|  |  |  | 3 DAI | NA |
|  |  |  | 4 DAI | NA |
|  |  |  | 5 DAI | NA |
|  | ROC22 | Susceptible | 0 DAI (Control) | - |
|  |  |  | 1 DAI | NA |
|  |  |  | 2 DAI | NA |
|  |  |  | 3 DAI | NA |
|  |  |  | 4 DAI | NA |
|  |  |  | 5 DAI | NA |

DAI: Days after inoculation. HAI: Hours after inoculation.

Read quality was assessed using FastQC software (Andrews, 2010) and visualized with MultiQC [54]. Quality control was performed using Trimmomatic [55] to remove adapter sequences and trim low-quality bases from both read ends (Phred score < 12), as well as internally using a sliding window approach (5 bp window; minimum average Phred score of 20). Reads shorter than 50% of their original length after trimming were discarded (<75bp for 150bp reads and <50bp for 100bp reads). Transcript quantification was carried out using Salmon version 1.1.0 [56], with reads aligned against the CDSs of the R570 cultivar genome [38].

### 2.6. Differential gene expression analyses

Transcript quantification data were imported into R using the tximport R package [57]. Differentially expressed genes (DEGs) were identified using DESeq2 R package version 3.9 [58]. Fold change (FC) values were obtained, and only DEGs with FDR-corrected *p*-values lower than 0.05 and log (|FC|) greater than 1 between the groups compared were considered significant. Differential gene expression analyses (DGE) were performed separately for each genotype within each study, comparing control and inoculated samples at different time points after infection. For the dataset including both resistant (YT93) and susceptible (ROC22) genotypes [22], comparisons between genotypes were also performed.

To compare DEG sets grouped according to smut resistance phenotype (resistant, intermediately resistant and susceptible), genes identified from comparisons between control and inoculated samples within the same genotype were considered. For each phenotype class, DEGs identified across different studies, genotypes and time points were combined by union into single gene sets. To characterize the biological processes associated with each resistance phenotype, Gene Ontology (GO) enrichment analyses were performed using the complete DEG set of each phenotype. Enrichment analyses were carried out using the topGO R package [59], and GO terms with a *p*-value lower than 0.05 were considered significantly enriched. The enriched GO terms were subsequently summarized and visualized using ReviGO version 1.8.2 [60], which was used to generate TreeMaps.

Venn diagrams were then generated using the ggVennDiagram R package [61] to compare the overlap among DEG sets grouped according to smut resistance phenotype. These comparisons were performed separately for up- and downregulated genes to identify phenotype-specific and shared transcriptional responses.

### 2.7. Co-expression network and functional enrichment analysis

A global weighted gene co-expression network was constructed using the WGCNA R package [62]. Gene quantification values were normalized using the transcripts per million (TPM) metric with the edgeR R package [63]. Pairwise Pearson correlations were calculated among TPM values for each gene, applying a power function fitted to scale-free topology. A soft-thresholding power (β) corresponding to an R² value of approximately 0.8 was estimated to approximate a scale-free topology model. Functional network modules were identified using the unweighted pair group method with arithmetic mean (UPGMA) based on a topological overlap matrix and dynamic tree cutting.

To further investigate the relationship between phenotype-specific DEG sets and the co-expression network modules, a Pearson’s chi-square test with Bonferroni correction (α = 0.05) was performed to assess the overrepresentation of DEG sets derived from the Venn diagrams within the network modules.

GO term enrichment was evaluated using the topGO R package [59] and conducted for the modules identified in the gene co-expression network, in order to characterize the associated biological processes. GO terms with a *p*-value lower than 0.05 were considered significantly enriched. The enriched GO terms obtained were then summarized and visualized using ReviGO version 1.8.2 [60], which was used to generate TreeMaps.

## 3. Results

### 3.1. Phenotypic data analyses

Resistance to smut was evaluated in 155 sugarcane genotypes, including commercial cultivars, clones, and species accessions, encompassing a range of previously characterized resistance levels to smut and representing diverse geographic origins (Table S1). Plants were subjected to bud inoculation with *S. scitamineum* and assessed in two independent assays conducted under greenhouse and field conditions. Three phenotypic variables were analysed: (i) percentage of whips emitted in greenhouse conditions, (ii) the AUDPC derived from this data, and (iii) percentage of whips emitted in field conditions. Analysis of the phenotypic data showed that it did not follow a normal distribution (Shapiro-Wilk test, *p* < 0.05); significant differences were detected among genotypes (Kruskal-Wallis test, *p* < 0.05), and no significant differences were observed among blocks (Kruskal-Wallis test, *p* > 0.05). Accordingly, for each phenotypic variable, block means were calculated, resulting in a final score per genotype for each of the three analyzed traits. Based on these values, scatter plots were generated and correlations among the three variables were estimated. Moderate, but significant correlations were observed between the percentage of whips emitted under field conditions and both AUDPC (R = 0.44, *p* < 0.001) and the percentage of whips emitted under greenhouse conditions (R = 0.397, *p* < 0.001). A high correlation was detected between AUDPC and the percentage of whips emitted under greenhouse conditions (R = 0.904, *p* < 0.001) (Fig. S1).

### 3.2. Genotyping and genetic analyses

Sequencing of the GBS library for 58 sugarcane genotypes generated 964,592,377 reads across two lanes. Both lanes showed high sequencing quality, with Phred scores above 32 across nearly all read positions, indicating a low base-calling error rate (Fig. S2). These data were combined with GBS data for another 97 genotypes generated previously, resulting in a dataset comprising 155 genotypes.

Variant calling was performed using the TASSEL4-POLY pipeline, and four reference sequences were used for alignment and SNP discovery: (i) the methyl-filtered genome of cultivar SP70-1143; (ii) the genome of the hybrid cultivar R570; (iii) the CDSs of this same genome; and (iv) the genome of the hybrid cultivar ZZ1. All analyses were conducted in parallel using the four references up to the final SNP filtering step. Among the references evaluated, the methyl-filtered SP70-1143 genome showed the best performance across the pipeline steps and filtering criteria, resulting in the highest single alignment rate and largest number of variants called. Therefore, this reference was selected for the subsequent step of genotype inference based on the proportion of observed allelic depths at each locus (Table 2).

**Table 2.** **Performance of the four genomic references in the SNP discovery and filtering pipeline**, including mapping rate, number of markers detected by TASSEL4-POLY and after filtering with VCFtools. The methyl-filtered reference, highlighted in bold, showed the best performance and was selected for subsequent analyses.

| **Reference** | **Mapping rate** |  | **Total markers** |  |
|---|---|---|---|---|
|  | **Total** | **Single** | **Tassel** | **VCFtools** |
| R570 genome | 95.99% | 3.44% | 39,811 | 1,719 |
| R570 CDSs | 58.51% | 8.58% | 117,472 | 13,545 |
| Methyl-filtered genome | 82.30% | 52.30% | 199,000 | 43,203 |
| ZZ1 genome | 96.70% | 2.12% | 21,592 | 576 |

Genotype inference was performed using ploidy levels of 6, 8, 10, 12, and 14. Among the evaluated ploidy levels, ploidy 12 yielded the highest number of high-quality SNP markers and was therefore selected, resulting in a final dataset of 33,335 SNP markers. These markers were then used to investigate the genetic structuring of the panel using two distinct approaches. A discriminant analysis of principal components (DAPC) indicated the presence of three subgroups in the panel (Figure 1a). In the case of principal component analysis (PCA) followed by K-means clustering, a variable number of clusters was obtained between methods used to identify cluster number: three groups according to the elbow method and two groups according to the silhouette method (Fig. S3). However, a high level of concordance (97%) was observed between the DAPC and the PCA with K-means clustering, with only four genotypes assigned to different groups by these two methods (Figure 1b). Overall, genotypes were largely segregated according to their classification: group 1 comprised 64% commercial varieties, group 2 was predominantly composed of IAC clones (85%), and group 3 consisted almost entirely of IAC clones (Table S2).

**Figure 1.**
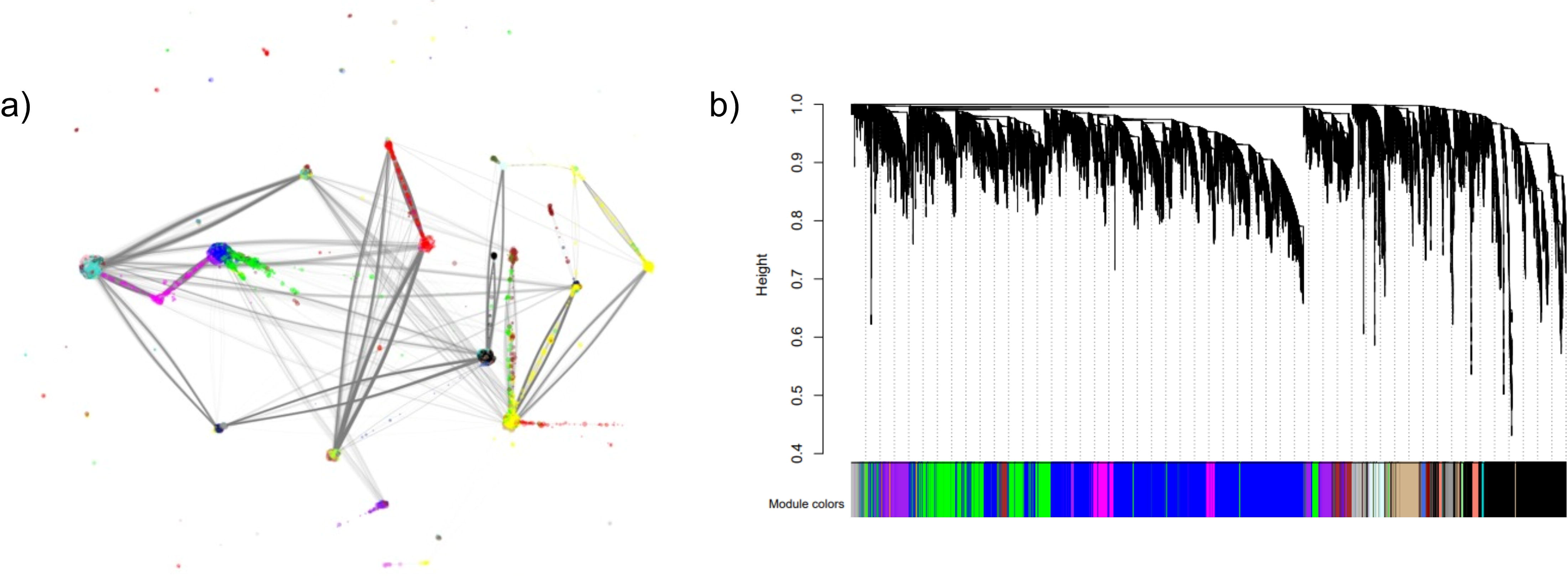
Genetic structure of the sugarcane genotype panel based on 33,335 SNP markers: (a) A discriminant analysis of principal components (DAPC), indicating the presence of three genetic subgroups; (b) Principal Component Analysis (PCA), showing the segregation of genotypes into three subgroups, colored according to the groups defined by DAPC.

### 3.3. Genome-wide association analyses

Based on the population structure analyses, different structure matrices (Q) were tested. For the PCA-based Q matrix, principal component scores from three PCs were used to represent population structure. For the DAPC-based Q matrix, the posterior membership probabilities of each genotype to the inferred clusters were used. Additionally, three distinct relationship (K) matrices, namely the realized kinship matrix, the complete autopolyploid matrix, and the pseudodiploid matrix, were calculated. Genome-wide association studies (GWASs) were performed with all possible combinations of Q + K matrices as factors and six marker-effect models (generalized, additive, reference simplex dominant, alternative simplex dominant, diploidized generalized, and diploidized additive) were tested. The best-performing parameter set consisted of the Q matrix derived from PCA combined with the pseudodiploid K matrix, as it yielded genomic inflation factor (λ) values closest to 1 (Fig. S4). Based on this combination, Q-Q plots were generated for all marker-effect models for the three traits under analysis (Figure 2). These plots enabled the evaluation of model adequacy and the exclusion of models that did not present appropriate λ inflation values. Accordingly, for the percentage of whip emission under greenhouse conditions and the AUDPC, the general model was excluded, whereas for the percentage of whip emission under field conditions, all models were retained for subsequent analyses.

**Figure 2.**
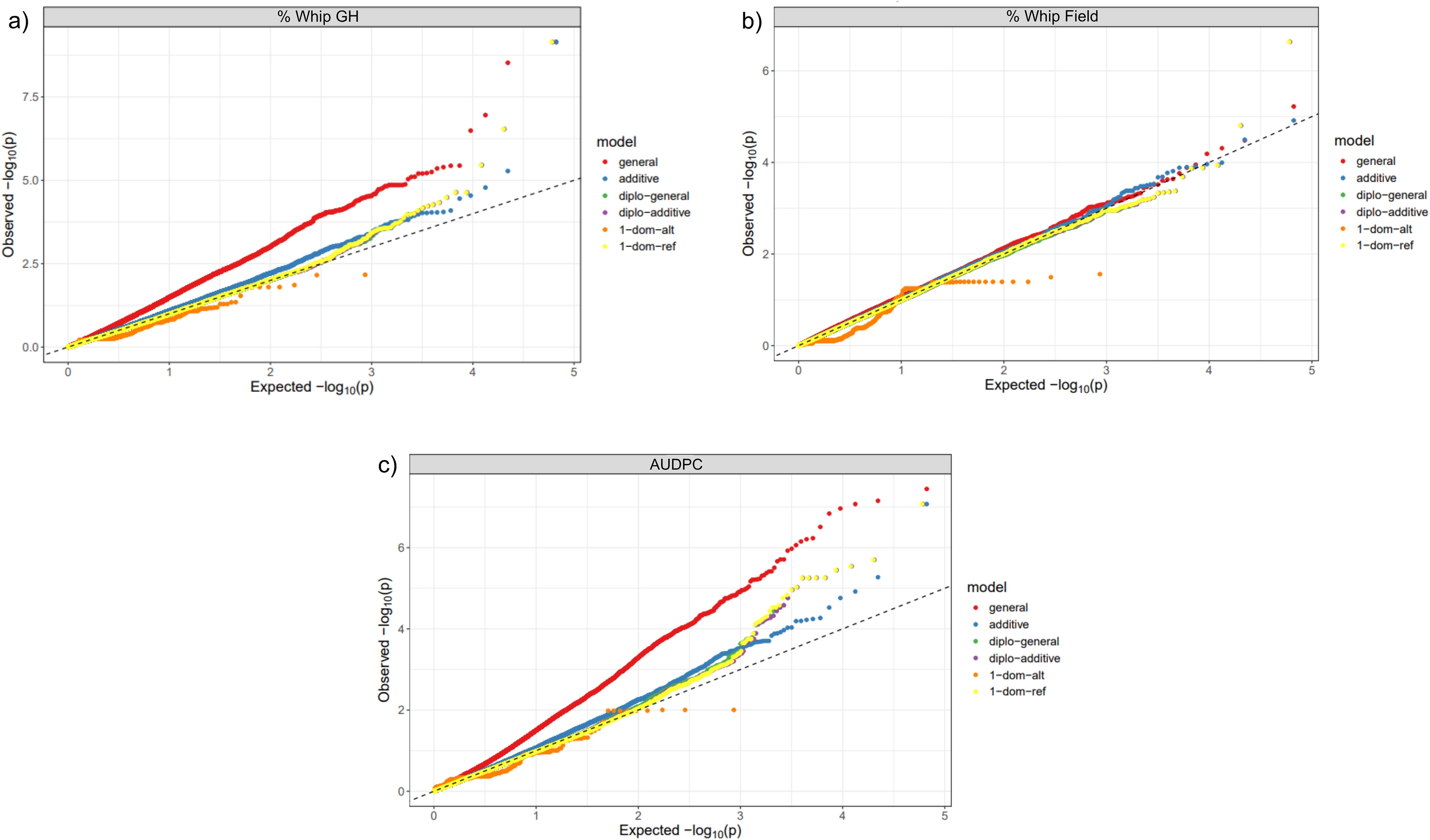
Quantile-quantile (QQ) plots of the different marker effect models tested for each trait: (a) percentage of whip emission under greenhouse conditions (% Whip GH), (b) percentage of whip emission under field conditions (% Whip Field), and (c) Area Under the Disease Progress Curve (AUDPC).

Markers significantly associated with smut resistance were assessed using the FDR correction, resulting in the identification of fourteen significant marker-trait associations (Table 3). The genomic positions of these markers were determined on the methyl-filtered genome, and their flanking sequences were aligned against the R570 genome. Based on the corresponding genomic regions, annotated genes were manually identified using the JBrowse genome browser available in Phytozome. One of the fourteen resistance-associated SNPs (16546) could not be successfully aligned to this reference.

**Table 3.** SNPs significantly associated with smut resistance in sugarcane identified by genome-wide association studies. The following information is provided: marker identifier, reference and alternative alleles, association significance (-log_10_(*p*)), estimated marker effect, and the GWAS marker-effect model(s) in which each SNP was identified as significant, for the three phenotypic traits: percentage of whip emission under greenhouse conditions (% Whip GH), percentage of whip emission under field conditions (% Whip Field), and Area Under the Disease Progress Curve (AUDPC).

| **SNP** | **Ref/alt allele** | **-log<sub>10</sub>(*p*)** | **Effect** | **Model(s)** |
|---|---|---|---|---|
| **% Whip GH** |  |  |  |  |
| 32075 | C/A | 9.14 | -35.71 | additive, diplo-general, diplo-additive, 1-dom-ref |
| 18183 | T/C | 6.54 | 34.88 | diplo-general, diplo-additive, 1-dom-ref |
| 4913 | G/T | 5.46 | 24.28 | diplo-general, diplo-additive, 1-dom-ref |
| 18475 | A/T | 2.17 | 23.87 | 1-dom-alt |
| **% Whip Field** |  |  |  |  |
| 3447 | G/A | 6.63 | -35.94 | diplo-general, diplo-additive, 1-dom-ref |
| **AUDPC** |  |  |  |  |
| 32075 | C/A | 7.07 | -763.73 | additive, diplo-general, diplo-additive, 1-dom-ref |
| 1676 | T/C | 4.76 | -760.57 | diplo-general, diplo-additive, 1-dom-ref |
| 4910 | G/A | 5.26 | 555.85 | diplo-general, diplo-additive, 1-dom-ref |
| 4912 | A/T | 5.26 | 555.85 | diplo-general, diplo-additive, 1-dom-ref |
| 9291 | A/G | 5.45 | 480.65 | diplo-general, diplo-additive, 1-dom-ref |
| 4913 | G/T | 5.70 | 577.30 | diplo-general, diplo-additive, 1-dom-ref |
| 16546 | C/G | 5.03 | 539.72 | diplo-general, diplo-additive, 1-dom-ref |
| 18183 | T/C | 5.54 | 740.98 | diplo-general, diplo-additive, 1-dom-ref |
| 20199 | A/G | 4.83 | 1222.15 | 1-dom-ref |
| 26739 | T/G | 5.25 | 556.98 | diplo-general, diplo-additive, 1-dom-ref |
| 26740 | C/G | 5.25 | 556.98 | diplo-general, diplo-additive, 1-dom-ref |
| 56 | G/T | 4.96 | 533.52 | diplo-general, diplo-additive, 1-dom-ref |

In cases where SNPs were located within genic regions, the corresponding gene annotations were retrieved. Two (32075 and 9291) of the thirteen resistance-associated SNPs that aligned successfully were not located within gene regions; therefore, for these cases, the nearest upstream and downstream genes were annotated (Table S3). For those located within genic regions, the reference and alternative nucleotide triplets were annotated to determine whether the substitution resulted in synonymous or non-synonymous amino acid changes. Of the ten SNPs located within gene regions, three resulted in non-synonymous mutations (Table S4).

### 3.4. Differential gene expression analyses

Reads generated in four studies investigating the transcriptomic response of five different sugarcane genotypes to inoculation with *S. scitamineum* were retrieved and processed. These included one smut-resistant, one tolerant, and three susceptible genotypes, sampled in various time points after inoculation (Table 1). Quality-filtered reads were used for gene expression quantification using the CDSs of the R570 cultivar genome as reference (Table S5).

Differential gene expression analyses were first performed between control and inoculated samples, revealing variation in the total number of DEGs across genotypes and post-inoculation time points (Table S6). As one of the datasets included both resistant and susceptible genotypes evaluated under the same experimental conditions, additional comparisons between these genotypes were also performed, considering the susceptible genotype as a reference. This allowed the assessment of expression differences associated with contrasting resistance phenotypes. The comparisons performed, the total numbers of DEGs identified in each of them, and their classification into up- and downregulated genes are presented in Table S7.

Based on these results, genes identified in the GWAS were cross-referenced with the sets of up- and downregulated genes derived from differential expression analyses between the resistant and susceptible genotypes, YT93 and ROC22, respectively, across five time points. Among the 16 resistance-associated genes identified, 11 were detected as differentially expressed in resistant genotypes compared with susceptible ones, exhibiting both up- and downregulation profiles across the evaluated conditions (Table 4, Table S7).

**Table 4.**
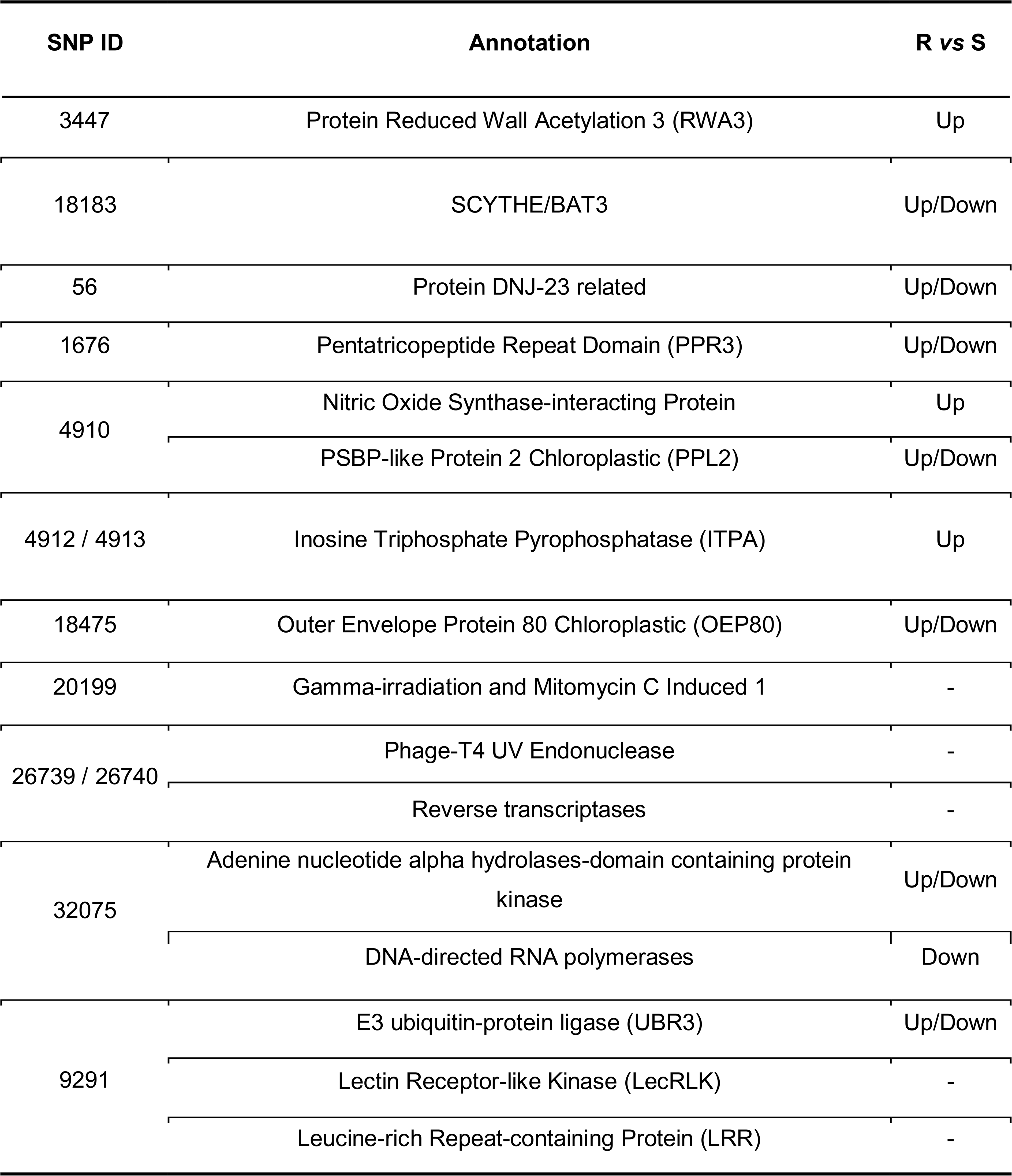
**Results of cross-referencing genes** identified in the GWAS and sets of genes up- and downregulated from differential expression analyses between resistant (YT93) and susceptible (ROC22) genotypes.

| **SNP ID** | **Annotation** | **R *vs* S** |
|---|---|---|
| 3447 | Protein Reduced Wall Acetylation 3 (RWA3) | Up |
| 18183 | SCYTHE/BAT3 | Up/Down |
| 56 | Protein DNJ-23 related | Up/Down |
| 1676 | Pentatricopeptide Repeat Domain (PPR3) | Up/Down |
| 4910 | Nitric Oxide Synthase-interacting Protein | Up |
|  | PSBP-like Protein 2 Chloroplastic (PPL2) | Up/Down |
| 4912 / 4913 | Inosine Triphosphate Pyrophosphatase (ITPA) | Up |
| 18475 | Outer Envelope Protein 80 Chloroplastic (OEP80) | Up/Down |
| 20199 | Gamma-irradiation and Mitomycin C Induced 1 | - |
| 26739 / 26740 | Phage-T4 UV Endonuclease | - |
|  | Reverse transcriptases | - |
| 32075 | Adenine nucleotide alpha hydrolases-domain containing protein kinase | Up/Down |
|  | DNA-directed RNA polymerases | Down |
| 9291 | E3 ubiquitin-protein ligase (UBR3) | Up/Down |
|  | Lectin Receptor-like Kinase (LecRLK) | - |
|  | Leucine-rich Repeat-containing Protein (LRR) | - |

In addition, a complementary analysis was performed to characterize transcriptional patterns associated with smut resistance phenotypic classification, based on comparisons between control and infected samples across different time points. DEG sets from genotypes within the same phenotypic class were integrated and further investigated through GO enrichment analyses and ReviGO-based functional summarization.

The functional summarization revealed distinct biological patterns associated with smut resistance phenotypic classification. Resistant genotypes showed a strong defense-related signature, with enrichment of processes associated with pathogen recognition, biotic stress responses, transcriptional regulation, and cell wall organization, suggesting activation of coordinated defense mechanisms (Fig. S5). Intermediately resistant genotypes displayed a predominantly structural and metabolic response, characterized by enrichment of cell wall remodeling and primary metabolic processes, indicating broader stress adaptation (Fig. S6). In contrast, susceptible genotypes exhibited mainly metabolic and cellular reorganization processes, including biosynthetic pathways and cellular organization, with limited representation of classical defense-related functions (Fig. S7).

To further investigate shared and phenotype-specific transcriptional responses, DEGs overlaps among resistance classes were visualized using Venn diagrams (Figure 3). Smut-resistant genotypes exhibited 6,239 exclusive upregulated and 3,034 exclusive downregulated genes, whereas intermediately resistant genotypes presented 753 and 648 exclusive up- and downregulated genes, respectively. Susceptible genotypes showed 4,350 exclusive upregulated and 3,714 exclusive downregulated genes. Additionally, a small set of genes displayed altered expression independently of the smut resistance level (85 upregulated and 8 downregulated), potentially representing general transcriptional responses to the pathogen (Figure 3).

**Figure 3.**
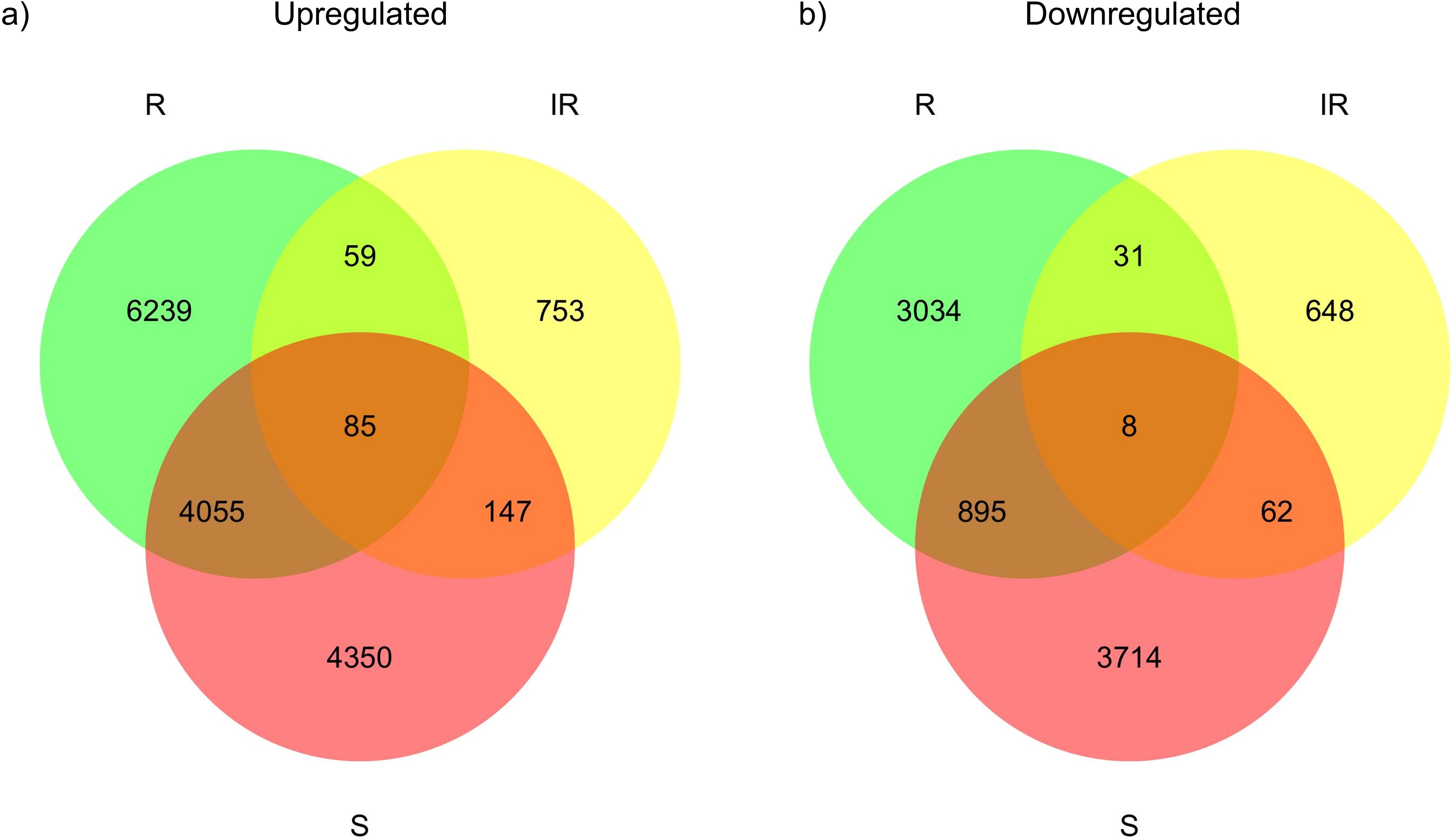
Venn diagrams showing the intersections among genes differentially expressed in response to *Sporisorium scitamineum* infection across resistant (R), intermediately resistant (IR), and susceptible (S) genotype groups. (a) Upregulated genes; (b) downregulated genes.

### 3.5. Gene coexpression network analyses

All 81 samples derived from the four selected studies were used to construct a clustering tree for outlier detection. Fourteen outlier samples were removed, with 67 samples retained for coexpression network construction (Fig. S8). The soft-thresholding power was set to β = 13, corresponding to the first point at which the scale-free topology fitting index reached approximately 0.8 (Fig. S9). 34 modules were identified in the resulting coexpression network, with sizes ranging from 13,617 (turquoise) to 155 (darkolivegreen) (Figure 4).

**Figure 4.**
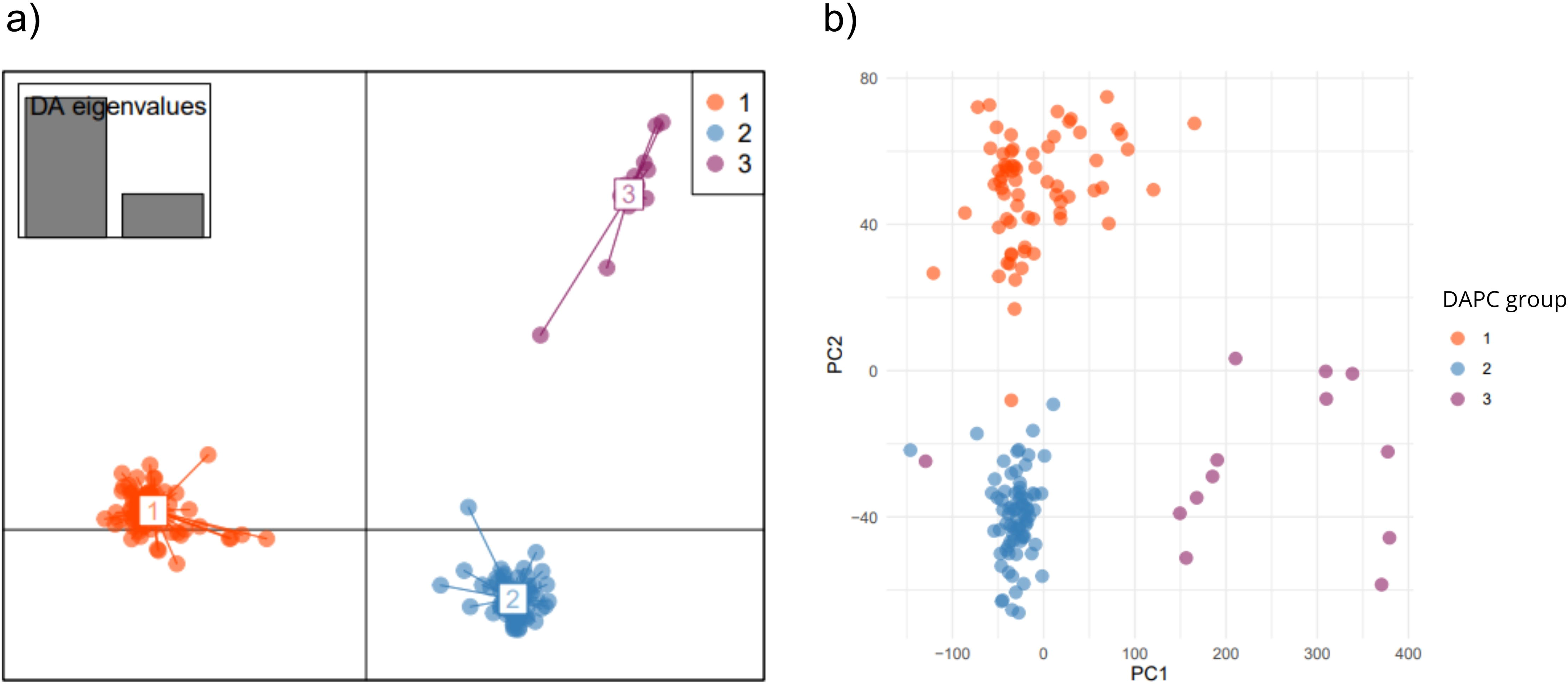
**Results of the gene coexpression analysis:** a) illustration of the coexpression network showing relationships among correlated genes; b) dendrogram highlighting modules and groups of genes clustered according to their expression patterns.

The identified modules were analysed for the overrepresentation of the DEGs identified between control and infected plants of smut-resistant, intermediately resistant, and susceptible sugarcane genotypes (Figure 3). DEGs exclusive to the resistant group were overrepresented in nine modules, among which three stood out due to their extremely high statistical significance: yellow (adjusted *p* value = 1.18×10^-99^), lightyellow (adjusted *p* value = 5.23×10^-174^), and skyblue (adjusted *p* value = 7.24×10^-170^). In comparison, four and ten modules were overrepresented with DEGs exclusive to the intermediately resistant and susceptible groups, respectively. Highly significant modules were also observed in these groups, including greenyellow (adjusted *p* value = 2.71×10^-56^) for intermediately resistant genotypes and black (adjusted *p* value = 4.21×10^-26^) and lightcyan (adjusted *p* value = 1.31×10^-26^) for susceptible genotypes (Table S8).

Functional enrichment summarization of the overrepresented modules for resistant and susceptible groups, visualized as TreeMaps, allowed the identification of key biological processes associated with resistance (Fig. S10) and susceptibility (Fig. S11) to sugarcane smut. In resistant genotypes, enriched processes included responses to wounding, phosphorylation, oxidative stress, photosynthesis, and cell wall organization and metabolism. Additionally, susceptible genotypes showed enrichment for processes related to regulation of molecular functions, autophagy, metabolic pathways, ion transport, and responses to oxidative stress and fungal infection.

Besides the overrepresentation analyses of DEGs from the different smut resistance phenotype groups, the modules identified in the co-expression network were important for identifying additional candidate genes associated with smut resistance. By cross-referencing the hub genes from the 34 co-expression modules with the complete set of DEGs obtained from the transcriptomic dataset (Table S7), 26 additional genes potentially involved in smut resistance were identified. Among these, 22 were differentially expressed in resistant genotypes compared with susceptible ones, being either up- or downregulated, representing particularly promising targets. The remaining four genes were differentially expressed only in comparisons among different days after infection within a same genotype (Table S9).

## 4. Discussion

This study aimed to investigate the genetic and transcriptional basis of sugarcane resistance to *S. scitamineum* through the integration of association mapping and transcriptomic analyses. A panel of 155 genotypes provided high-quality genotypic and phenotypic data, supporting the robustness and reliability of the subsequent analyses. Additionally, publicly available transcriptomic datasets of sugarcane infected with smut were processed and analyzed, adding an important complementary layer of evidence that enabled the integration of GWAS-identified candidate genes with differential expression and co-expression network analyses.

### Genome complexity and bioinformatic analyses

The genomic complexity of sugarcane may still represent a challenge for association studies. Its highly polyploid and aneuploid genome can hinder read mapping, SNP discovery, and genotype inference, often requiring simplifications into diploid-based analytical frameworks [1, 3, 4]. This limitation was reflected in the superior performance of the methyl-filtered SP70-1143 reference genome, whose reduced repetitive content favored unique read alignment and yielded a greater number of high-confidence variants than the chromosome-scale references evaluated. These results suggest that current analytical pipelines may not yet fully exploit the information available in complete sugarcane genome assemblies.

To address part of this complexity, genotype inference in this study incorporated allele dosage estimates based on allelic read-depth proportions rather than relying exclusively on discrete diploid genotype classes. This strategy allows a more realistic representation of the multiple allele copies present at each locus and may improve the detection of marker-trait associations in polyploid species [40].

Beyond these methodological considerations, the integration of genomic and transcriptomic datasets provided a comprehensive framework for investigating resistance-associated loci, expression patterns, and co-expression network. Such approaches may contribute to a deeper understanding of the sugarcane-smut interaction and support the development of molecular tools for resistance breeding. The following sections examine how these complementary lines of evidence contribute to our understanding of the molecular architecture of resistance to *S. scitamineum* in sugarcane.

#### Key genes and process involved in resistance to smut

The resistance-associated SNPs identified in this study, together with their linked genes, expand the current understanding of the molecular basis underlying sugarcane responses to smut infection. Functional annotation of these candidates revealed genes involved in key defense-related processes, including cell wall remodeling, pathogen perception and signaling, redox homeostasis, genome maintenance, RNA-mediated regulation, and chloroplast protein homeostasis. Collectively, these findings suggest that resistance to *S. scitamineum* depends on the activation of multiple molecular pathways that contribute to pathogen restriction and cellular adaptation during infection. The main biological processes involved in smut resistance and the candidate genes associated with these processes are discussed below, based on the evidence obtained in this study and the available literature.

### Cell wall modification

Reduced Wall Acetylation (RWA) proteins, such as RWA3, were the first class of proteins identified as regulators of plant cell wall polysaccharide acetylation [64]. In *Arabidopsis*, RWA genes are regulated by the secondary cell wall master regulator SND1, are highly expressed during secondary wall formation, and are required for proper wall thickening and cell differentiation [65]. Functional studies have shown that simultaneous knockout of the four RWA genes substantially reduces cell wall acetylation, including xylan O-acetylation, impairing secondary wall development [65,66,67]. Although reduced wall O-acetylation has been associated with increased fungal resistance in other pathosystems [64,68], our results suggest that sugarcane resistance to *S. scitamineum* is associated with increased cell wall O-acetylation, as supported by the upregulation of RWA3 in resistant genotypes. Notably, RWA3 was also supported by the GWAS analysis, providing independent genetic evidence for its potential role in resistance (Table 4). The identification of RWA3 by both genomic and transcriptomic approaches considerably strengthens its candidacy, placing it among the most compelling resistance-associated genes uncovered in this study. These results further suggest that cell wall O-acetylation may play a previously underappreciated role in sugarcane resistance to smut, revealing a potentially distinct defense strategy in this pathosystem.

Because increased wall acetylation also contributes to biomass recalcitrance by limiting enzymatic hydrolysis and reducing saccharification efficiency, differential RWA expression directly affects biomass processability [67,69,70]. The association between higher RWA3 expression, increased cell wall acetylation, and biomass recalcitrance also suggests a potential trade-off between cell wall properties associated with biomass utilization and resistance to smut. Such a trade-off could represent an important challenge for breeding programs aiming to combine high biomass productivity with durable disease resistance [1,10].

### Pathogen perception and defense signaling

The role of the plant cell wall in resistance extends beyond structural genes and includes key regulators of pathogen perception and defense signaling, which act during the early stages of plant immunity. Plant immune responses are generally organized into two major layers: PAMP-triggered immunity (PTI), initiated by recognition of microbe- and pathogen-associated molecular patterns (MAMPs/PAMPs) through plasma membrane pattern recognition receptors (PRRs), and effector-triggered immunity (ETI), activated by intracellular recognition of pathogen effectors. PTI activates defense responses such as MAPK cascades, calcium influx, and ROS bursts, whereas ETI induces stronger responses, including hypersensitive cell death and systemic acquired resistance [71,72,73,74].

Among PRRs, lectin receptor-like kinases (LecRLKs) are widely associated with biotic stress responses due to their ability to perceive extracellular signals and activate downstream defense pathways [73,75]. In our study, a genomic region significantly associated with smut resistance was found in proximity to a LecRLK-encoding gene (Table S3), suggesting a potential role for receptor-mediated pathogen recognition in the sugarcane response to *S. scitamineum*. Their role in fungal resistance is well established, as exemplified by the rice G-type LecRLK Pi-d2, in which a single amino acid substitution is sufficient to confer resistance to *Magnaporthe grisea* in rice, and L- and G-type LecRLKs associated with differential resistance to *Sphaerulina musiva* in *Populus trichocarpa* [73,76,77]. Collectively, these findings reinforce the importance of LecRLKs in sugarcane resistance to smut, supporting our results and underscoring the conservation of receptor-mediated fungal recognition mechanisms across plant species.

Likewise, leucine-rich repeat (LRR)-containing proteins, including LRR-RLKs and NBS-LRR receptors, play central roles in pathogen recognition and downstream defense activation [78,79]. In this study, LRR- and RLK-related sequences were identified near SNP 9291, whose alternative allele was associated with increased AUDPC and greater smut susceptibility relative to the reference allele (Tables 3 and S3). Although this SNP is intergenic and unlikely to directly alter protein structure, it may affect transcriptional regulation or be linked with other polymorphisms of nearby defense-related genes. This interpretation is supported by previous sugarcane studies identifying LRR- and RLK-related genes associated with resistance loci and as hub genes in co-expression networks [22,80].

Consistently, homology analysis revealed 72.9% similarity between the LRR gene identified here and an NBS-LRR disease resistance protein from *Oryza sativa*. NBS-LRR genes are broadly associated with enhanced resistance across crops and have been successfully used in breeding strategies, such as AhRRS5-mediated resistance to *Ralstonia solanacearum* in transgenic tobacco [81]. Together, these findings strengthen the evidence that LRR- and RLK-related genes contribute to smut resistance and position them as promising targets for molecular breeding strategies for disease resistance improvement.

### Oxidative stress and DNA damage responses

Activation of PTI and ETI is accompanied by an oxidative burst, characterized by increased ROS and NO production [74]. In this study, the Nitric Oxide Synthase-interacting protein gene was associated with smut resistance through a SNP in the 3′UTR region and was upregulated in resistant genotypes, suggesting enhanced NO signaling and redox-mediated defense responses during the sugarcane-smut interaction (Tables 4 and S4).

Because excessive ROS accumulation can damage nucleic acids directly or through reactive byproducts of lipid peroxidation, plants activate DNA damage response pathways to preserve genome integrity, whereas failure to properly resolve these lesions may result in mutations, metabolic dysfunctions, or cell death, with proteins such as SCYTHE/BAT3 contributing to apoptosis regulation [82,83,84]. Consistently, genes involved in DNA repair and genome stability, including Gamma-irradiation and Mitomycin C Induced 1, Phage-T4 UV Endonuclease, reverse transcriptases, and DNA-directed RNA polymerases, were associated with smut resistance, reinforcing the importance of efficient DNA repair for stress adaptation and cellular survival (Table 4) [83,84,85].

Notably, SNPs in Gamma-irradiation and Mitomycin C Induced 1 (SNP 20199), Phage-T4 UV endonuclease, and reverse transcriptases (SNPs 26739/26740) resulted in non-synonymous substitutions associated with increased susceptibility. These changes replaced histidine or leucine with arginine, introducing greater positive charge that may alter protein structure and DNA-binding affinity (Tables 3 and S4). Such structural changes are likely to impair DNA repair efficiency under pathogen-induced oxidative stress, compromising genome maintenance and thereby contributing to increased susceptibility to infection.

### Genome integrity and RNA-mediated regulation

Building on the effects of oxidative stress on DNA integrity, plants activate additional protective mechanisms to preserve genome stability under stress. The Adenine nucleotide alpha hydrolase-domain containing protein kinase encodes a protein harboring both a universal stress protein (USP)-like domain and a kinase domain. In plants, USPs contribute to protein stability, prevention of macromolecular damage, and maintenance of cellular homeostasis under stress [86,87,88]. The identification of this gene through the GWAS and transcriptomic analysis suggests its potential involvement in sugarcane resistance to smut (Table 4). The presence of a kinase domain further suggests a role in integrating stress perception with downstream adaptive signaling, potentially contributing to sugarcane resistance to smut.

Similarly, maintenance of nucleotide pool integrity is essential under oxidative stress. Inosine Triphosphate Pyrophosphatase (ITPA) prevents the incorporation of damaged purine nucleotides into RNA and DNA by hydrolyzing noncanonical nucleotides, thereby limiting mutagenesis and genomic instability [89,90,91]. The higher ITPA expression observed in resistant genotypes suggests enhanced nucleotide sanitization capacity, which likely contributes to improved genome stability under smut-induced oxidative stress, thereby reinforcing cellular homeostasis during infection (Table 4).

RNA integrity is also influenced by Pentatricopeptide Repeat Domain (PPR3), whose proteins regulate organellar RNA processing, stability, and translation, being crucial for plant growth and development, including cytoplasmic male sterility, seed development, photosynthesis, and responses to biotic and abiotic stresses [92,93,94]. Defects in these proteins often lead to organellar dysfunction and ROS accumulation through impaired post-transcriptional regulation [94]. In this study, a non-synonymous substitution in PPR3 showed a negative SNP effect associated with resistance, suggesting that this variant may optimize RNA-mediated regulatory processes involved in stress adaptation (Tables 3 and S4).

### Chloroplast function and protein homeostasis

Chloroplasts act as dynamic environmental sensors that coordinate stress adaptation through bidirectional signaling with the nucleus, mediated in part by signaling molecules such as ROS and NO [95,96]. In this context, PSBP-like Protein 2 (PPL2) and Outer Envelope Protein 80 (OEP80) emerged as relevant candidates for sugarcane resistance to smut (Table 4). PPL2, an extrinsic photosystem II subunit, is associated with photosynthetic performance under stress, whereas OEP80 contributes to chloroplast membrane assembly, permeability, protein translocation, and intercompartmental communication, all of which are essential for chloroplast functionality [97,98,99,100,101].

Maintenance of chloroplast function during smut infection also depends on protein quality control systems, including chloroplast heat shock proteins that refold or degrade damaged proteins [102,103]. Consistently, the DNJ-23-related gene identified here, encoding a DNAJ/HSP40 molecular chaperone, likely contributes to chloroplast protein homeostasis by stabilizing or removing proteins damaged under oxidative stress (Table 4) [103,104].

Protein homeostasis is further regulated by post-translational modifications, particularly ubiquitination, which controls protein stability and turnover through the ubiquitin-proteasome system [105,106]. The UBR3 E3 ubiquitin ligase identified in this study may contribute to immune regulation by modulating the abundance of defense-related proteins involved in both PTI and ETI (Table 4). This is consistent with the previous sugarcane GWAS study linking ubiquitin-proteasome components to smut resistance [80]. Together, these findings highlight the importance of chloroplast integrity, protein quality control, and post-translational regulation in coordinating effective defense responses against smut infection.

#### Co-expression modules associated with distinct resistance levels

Transcriptomic analyses provided a complementary layer to the genomic results, revealing that sugarcane genotypes with contrasting smut resistance phenotypes differ not only in the genes they regulate but also in the organization of their transcriptional responses. Functional enrichment of phenotype-associated DEGs identified distinct biological signatures across resistance classes, whereas the identification of phenotype-specific DEGs refined these signatures by distinguishing genes unique to each resistance class from those shared across phenotypes. Co-expression network analysis of these exclusive gene sets further demonstrated that resistant, intermediately resistant, and susceptible genotypes are characterized by distinct transcriptional modules, supporting the view that the molecular response to *S. scitamineum* is coordinated in a phenotype-dependent manner (Table S8).

Our analyses identified nine co-expression modules with a significant overrepresentation of DEGs exclusive to smut-resistant sugarcane genotypes. With the exception of the steelblue module, these modules did not overlap with those associated with DEGs exclusive to smut-susceptible genotypes, suggesting that fungal infection triggers distinct transcriptional programs depending on the resistance level, and that these responses are directly associated with the plant’s ability to withstand the pathogen. Intermediately resistant genotypes, in turn, displayed a mixed pattern, with significant modules overlapping both resistant (e.g., greenyellow and skyblue) and susceptible (e.g., salmon) groups, indicating that their response combines elements of both resistance and susceptibility.

Modules with an overrepresentation of DEGs exclusive to smut-resistant sugarcane genotypes were enriched for processes related to wounding response, phosphorylation, oxidative stress, photosynthesis, and cell wall organization and biogenesis. The recurrent enrichment of cell wall-related processes across highly significant modules reinforces the importance of cell wall remodeling as a structural barrier against pathogen invasion. Enrichment of amino sugar metabolism, amino acid transport, and carboxylic acid metabolism further suggests extensive metabolic reprogramming, whereas the prevalence of phosphorylation highlights the central role of signaling cascades in coordinating defense responses.

In contrast, modules with an overrepresentation of DEGs exclusive to smut-susceptible genotypes were enriched for negative regulation of molecular functions, autophagy, and stress-related responses. Additional enrichment of hormone regulation, trehalose biosynthesis, ion transport, and lipid metabolism reflects metabolic adjustment and damage mitigation. Although defense-related processes, including response to fungus and oxidative stress, were also enriched, these patterns likely reflect stress adaptation rather than effective resistance mechanisms capable of restricting pathogen progression.

#### Central hub genes in transcriptional responses to smut

In addition to enabling the identification of co-expression modules associated with distinct resistance levels, co-expression network analysis also provides insights into key regulatory genes involved in plant responses to pathogen infection, represented by hub genes. Due to their high connectivity within modules, these genes are considered central regulatory elements and are often strongly associated with traits of interest [62]. Among all pathogen-responsive DEGs identified across the analyzed transcriptomic datasets, 26 were classified as hub genes, some of which deserve particular attention (Table S9).

Similar to RWA3, identified in the GWAS, the cell wall-related gene Glycine-rich Cell Wall Structural Protein was also upregulated in resistant genotypes. Glycine-rich proteins are associated with stress signaling and antimicrobial responses, and their overexpression has been linked to enhanced tolerance to both abiotic and biotic stresses [107], supporting a multi-omic signal for the role of cell wall-related processes in sugarcane resistance to smut.

Other upregulated hub genes in resistant genotypes included Protein kinase - Gnk2-homologous domain and HVA22-like Protein E. Gnk2 homologs exhibit antifungal activity and have been associated with increased tolerance to *Phytophthora cinnamomi* in transgenic plants [108,109]. In the same way, HVA22 family genes, known for their roles in stress adaptation, have been associated with enhanced resistance to blast disease in rice [110] and with tolerance to abiotic stress in several plant species, including *Brassica napus* [111], pineapple [112], soybean [113], cotton [114], tomato [115], and citrus [116].

Additional evidence comes from seven hub DEGs belonging to modules overrepresented in resistant genotypes: Cysteine-rich receptor-like protein kinase 25 (lightyellow), HVA22-like protein E (skyblue), 3-ketoacyl-COA synthase 5-related (steelblue), Sulfate transporter 3.1-related (white), Chaperone protein DNAJ A6 Chloroplastic (yellow), Flavonoid 3’-monooxygenase (greenyellow) and ABC Transporter domain-containing protein (saddlebrown). Their differential expression, combined with their identification as central regulators in resistance-associated modules, provides robust multi-level evidence supporting their involvement in smut resistance.

Further support emerged from integrating these findings with GWAS data. Considering the approximately two million base pair linkage disequilibrium windows reported for sugarcane [26], expansion of the annotation window around resistance-associated SNPs identified two additional hub genes: Large subunit ribosomal protein L7Ae (RP-L7Ae), upregulated in resistant plants, and Oxysterol-binding Protein-related Protein 3C, downregulated in resistant plants.

Together, the integration of differential expression, co-expression network structure, and genomic association provides convergent multi-omic evidence supporting these genes as high-confidence candidates associated with sugarcane resistance to smut, positioning them as targets for functional validation and resistance-oriented breeding strategies.

## 5. Conclusions

Collectively, the main findings of this study support a multilayered model of sugarcane resistance to *S. scitamineum*, involving the coordinated regulation of genes associated in structural reinforcement, immune signaling, redox homeostasis, genome maintenance, and cellular adaptation mechanisms. Importantly, the integration of genomic and transcriptomic analyses provided a robust multi-omic framework that enabled the convergence of independent lines of evidence, strengthening the identification and prioritization of resistance-associated genes. This integrative approach not only enhances confidence in the candidate genes and molecular markers identified, but also offers a more comprehensive and systems-level view of the sugarcane-smut interaction. Furthermore, the identified candidate genes and resistance-associated markers provide promising molecular resources for functional validation and offer strategic targets for breeding programs aimed at developing sugarcane cultivars with enhanced resistance to smut.

## Acknowledgements

The authors are grateful to the São Paulo Research Foundation (FAPESP, 2024/14309-8, 2022/04006-2 and 17/25258-1) and Coordination for the Improvement of Higher Education Personnel (CAPES, 88887.994147/2024-00) for supporting the project and researchers.

## Data availability

The raw sequencing data used in this article have been submitted to the SRA under BioProjects PRJNA702641, PRJNA1416341, and PRJNA1557411. Table S6 is available on FigShare under the DOI 10.6084/m9.figshare.34918068. Figures S10 and S11 are available at high resolution on FigShare under the DOIs 10.6084/m9.figshare.34918326 and 10.6084/m9.figshare.34918341, respectively.

## CRediT

D.T.: Investigation, Data curation, Formal analysis, Visualization, Writing - original draft, Writing - review & editing. R.J.G.P.: Conceptualization, Methodology, Investigation, Data curation, Formal analysis, Writing - review & editing. L.R.P.: Conceptualization, Methodology, Supervision, Writing - review & editing. A.P.S.: Conceptualization, Methodology, Supervision, Writing - review & editing. L.F.S.: Investigation. L.O.S.A.: Investigation. I.A.A.: Investigation. F.B.S.: Investigation. A.C.L.M.: Investigation, Writing - review & editing. D.A.S.: Writing - review & editing.

All authors read and approved the final manuscript.

## Declaration/Statements

The authors declare that they have no known competing financial interests or personal relationships that could have appeared to influence the work reported in this paper.

