## Supplementary Material for "Multi-omics integration reveals candidate genes and molecular mechanisms associated with resistance to sugarcane smut"

^d^ Universidade Estadual Paulista “Júlio de Mesquita Filho”, Campus de Jaboticabal, SP.

^e^ Centro de Biologia Molecular e Engenharia Genética (CBMEG), Universidade Estadual de Campinas (UNICAMP), Campinas, SP, Brasil.

^f^ Instituto Agronômico Campinas (IAC), Divisão Avançada de Pesquisa e Desenvolvimento de Cana, Ribeirão Preto, SP.

**Table S1. List of the 155 sugarcane genotypes presented in the association panel**, including commercial cultivars, breeding program clones, and *Saccharum* species accessions. The table presents genotype classification (A = Instituto Agronômico de Campinas (IAC) clone; B = commercial cultivar; C = species accession), as well as female and male parental information when available.

| **Genotype** | **A = IAC clone;**  **B = commercial;**  **C = species** | **Female parent** | **Male parent** |
| --- | --- | --- | --- |
| Alho |  | *S. officinarum* | Unknown |
| CO213 | B | COLK 8102 | CO775 |
| CO419 | B | POJ 2878 | CO290 |
| CTC07 | B | RTT8618 | Policruzamento |
| CTC08 | B | SP813492 | Policruzamento |
| CTC12 | B | SP801842 | SP811747 |
| CTC15 | B | SP842025 | Policruzamento |
| CTC21 | B | CT971246 | SP832847 |
| CTC22 | B | SP803280 | Policruzamento |
| IAC-C01 | A | IAC133247 | IJ76482 |
| IAC-C02 | A | IAC127060 | IACSP04065 |
| IAC-C03 | A | IAC127059 | IACSP04065 |
| IAC-C04 | A | CTC9 | IAC133172 |
| IAC-C05 | A | IAC133176 | IACSP974039 |
| IAC-C06 | A | IAC133247 | IACSP994010 |
| IAC-C07 | A | IACSP013127 | IACSP962008 |
| IAC-C08 | A | IAC133235 | IACSP018158 |
| IAC-C09 | A | IAC133435 | IACSP976628 |
| IAC-C10 | A | IAC133264 | IACSP976619 |
| IAC-C11 | A | IACSP047070 | IAC127064 |
| IAC-C12 | A | Unknown | Unknown |
| IAC-C13 | A | IACSP976680 | IAC127064 |
| IAC-C14 | A | CO419 | C0356 |
| IAC-C15 | A | Unknown | Unknown |
| IAC-C16 | A | US71399 | Policruzamento |
| IAC873396 | B | CO740 | SP701143 |
| IAC-C17 | A | RB785148 | Policruzamento |
| IAC-C18 | A | SP813137 | Policruzamento |
| IAC-C19 | A | RB855465 | GLAGAH |
| IAC-C20 | A | RB855465 | GLAGAH |
| IAC-C21 | A | NG26011 | Policruzamento |
| IAC-C22 | A | NG26011 | Policruzamento |
| IAC-C23 | A | SES069 | Policruzamento |
| IAC-C24 | A | SES069 | Policruzamento |
| IAC-C25 | A | SES069 | Policruzamento |
| IAC-C26 | A | SES069 | Policruzamento |
| IAC-C27 | A | SES069 | Policruzamento |
| IAC-C28 | A | IAC915155 | CT961414 |
| IAC-C29 | A | SP911049 | CT931231 |
| IAC-C30 | A | IACSP955000 | CT961414 |
| IAC-C31 | A | IACSP953028 | CT961414 |
| IAC-C32 | A | IACSP955011 | CT961414 |
| IAC-C33 | A | White transparent | SP80-3280 |
| IAC-C34 | A | IACSP972053 | SP88721 |
| IAC-C35 | A | IACSP972053 | RB85453 |
| IAC-C36 | A | IACSP972053 | RB855453 |
| IAC-C37 | A | CT943165 | IAC911099 |
| IAC-C38 | A | IACSP962042 | CT014455 |
| IAC-C39 | A | IACSP955000 | CT961307 |
| IAC-C40 | A | IACSP955000 | RB72454 |
| IAC-C41 | A | IACSP963069 | IACSP952078 |
| IAC-C42 | A | IACSP963076 | CTC4 |
| IAC-C43 | A | R570 | IAC873396 |
| IAC-C44 | A | RB835486 | SP775181 |
| IAC-C45 | A | RB835486 | SP775181 |
| IAC-C46 | A | IAC912195 | SELF |
| IACSP013127 | B | RB835486 | SP775181 |
| IAC-C47 | A | SP913059 | CTC957 |
| IACSP015503 | B | SP913059 | CTC957 |
| IAC-C48 | A | RB835486 | SP775181 |
| IAC-C49 | A | IAC913093 | SP87432 |
| IAC-C50 | A | R570 | IAC873396 |
| IAC-C51 | A | IAC913093 | SP87365 |
| IAC-C52 | A | RB835486 | SP775181 |
| IAC-C53 | A | SP973060 | SP701143 |
| IAC-C54 | A | SP901161 | SP901644 |
| IAC-C55 | A | SP924230 | SP88813 |
| IAC-C56 | A | IAC914216 | SP803280 |
| IAC-C57 | A | PO8862 | CTC9 |
| IAC-C58 | A | IACSP955037 | Policruzamento |
| IAC-C59 | A | SP913059 | IACSP972077 |
| IAC-C60 | A | SP913059 | IACSP973391 |
| IAC-C61 | A | IAC913093 | CTC9638 |
| IAC-C62 | A | IAC912288 | SP80185 |
| IAC-C63 | A | IAC912288 | SP963092 |
| IAC-C64 | A | IACSP953028 | SP832847 |
| IAC-C65 | A | IACSP953018 | IACSP933046 |
| IAC-C66 | A | IACSP953018 | IACSP933046 |
| IAC-C67 | A | IACSP966026 | RB855453 |
| IAC-C68 | A | IACSP956114 | IACSP953050 |
| IAC-C69 | A | IACSP966026 | RB855453 |
| IAC-C70 | A | PO8862 | CTC9 |
| IAC-C71 | A | IACSP953028 | SP775181 |
| IAC-C72 | A | SP913059 | IACSP966026 |
| IAC-C73 | A | IACSP956114 | IACSP933046 |
| IAC-C74 | A | IACSP953018 | SP963092 |
| IAC-C75 | A | IACSP953018 | SP963092 |
| IAC-C76 | A | IACSP953028 | SP924221 |
| IAC-C77 | A | IACSP972109 | IACSP956114 |
| IAC-C78 | A | IAC911099 | SP931124 |
| IACSP932060 | B | RB765418 | Policruzamento |
| IACSP933046 | B | SP791011 | Policruzamento |
| IAC-C79 | A | SP791011 | Policruzamento |
| IACSP942094 | B | SP847017 | Policruzamento |
| IACSP942101 | B | SP775181 | RB785148 |
| IACSP944004 | B | SP26108 | SP775181 |
| IAC-C80 | A | SP842189 | SP801842 |
| IAC-C81 | A | CTC4 | IACSP933046 |
| IACSP955000 | B | SP842066 | SP80185 |
| IACSP955037 | A | SP842066 | IAC873396 |
| IACSP955094 | B | SP803280 | Policruzamento |
| IAC-C82 | A | IAC873187 | CTC9019 |
| IAC-C83 | A | SP80144 | Policruzamento |
| IACSP962042 | B | SP815193 | SP775181 |
| IAC-C84 | A | SP826108 | Policruzamento |
| IAC-C85 | A | SP803280 | Policruzamento |
| IAC-C86 | A | SP847017 | Policruzamento |
| IAC-C87 | A | RB855035 | SP80185 |
| IAC-C88 | A | RB855113 | RB855453 |
| IAC-C89 | A | RB835486 | RB855453 |
| IAC-C90 | A | RB835486 | SELF |
| IAC-C91 | A | SP842066 | SP847017 |
| IAC-C92 | A | IAC913093 | SP823530 |
| IAC-C93 | A | SP813251 | SP823697 |
| IAC-C94 | A | SP851682 | SP80185 |
| IAC-C95 | A | CP701133 | SP832847 |
| IAC-C96 | A | SP921584 | Unknown |
| IAC-C97 | A | SP901616 | Policruzamento |
| IAC-C98 | A | SP842025 | RB855113 |
| IAC-C99 | C | *S. robustum* | - |
| IAC-C100 | C | *S. spontaneum* | - |
| IAC-C101 | C | *S. spontaneum* | - |
| IAC-C102 | C | *S. spontaneum* | - |
| IAC-C103 | C | *S. officinarum* | - |
| KASSOER | B | *S. officinarum* | GLAGAH |
| KRAKATAU | C | *S. spontaneum* | Unknown |
| NA5679 | B | CO419 | CO419 |
| NCO310 | B | CO421 | CO312 |
| NG5712 | C | *S. robustum* | Unknown |
| NG57213 | C | Unknown | Unknown |
| POJ2878 | B | POJ2364 | EK28 |
| R570 | B | H393633 | M20246 |
| RAGNAR | B | CO270 | MQ3371 |
| RB855156 | B | RB72454 | TUC717 |
| RB855453 | B | TUC717 | Unknown |
| RB867515 | B | RB72454 | Unknown |
| RB92579 | B | RB755126 | RB72199 |
| RB935744 | B | RB835089 | RB765418 |
| SES205A | B | *S.spontaneum* | Unknown |
| SP716163 | B | NA5679 | Unknown |
| SP701143 | B | IAC4865 | Unknown |
| SP80185 | B | BO17 | Unknown |
| SP803280 | B | SP711088 | H575028 |
| SP813250 | B | CP701547 | SP711279 |
| SP832847 | B | HJ5741 | SP701143 |
| SP835073 | B | SP711406 | SP711088 |
| SP901638 | B | SP784601 | Unknown |
| SP903414 | B | SP801079 | SP823544 |
| SP911049 | B | SP803328 | SP813250 |
| U55714105 | B | *S. spontanuem* | Unknown |
| US571415 | C | Unknown | Unknown |
| VERTIX2 | B | IAC911099 | KRAKATAU |
| VERTIX3 | B | IACBIO79 | Unknown |
| VERTIX7 | B | IACBIO123 | Unknown |
| VERTIX9 | B | IACSP963055 | KRAKATAU |

**Table S2. Group assignment of the 155 sugarcane genotypes in the association panel** based on discriminant analysis of principal components (DAPC) and principal component analysis (PCA) followed by K-means clustering. The table presents the cluster classification of each genotype according to both methods.

| **Genotype** | **Group** | |
| --- | --- | --- |
|  | **DAPC** | **PCA** |
| Alho | 2 | 2 |
| CO213 | 1 | 1 |
| CO419 | 1 | 1 |
| CTC07 | 1 | 1 |
| CTC08 | 1 | 1 |
| CTC12 | 1 | 1 |
| CTC15 | 2 | 2 |
| CTC21 | 1 | 1 |
| CTC22 | 1 | 1 |
| IAC-C01 | 1 | 1 |
| IAC-C02 | 1 | 1 |
| IAC-C03 | 1 | 1 |
| IAC-C04 | 1 | 1 |
| IAC-C05 | 1 | 1 |
| IAC-C06 | 1 | 1 |
| IAC-C07 | 1 | 1 |
| IAC-C08 | 1 | 1 |
| IAC-C09 | 1 | 1 |
| IAC-C10 | 1 | 1 |
| IAC-C11 | 1 | 1 |
| IAC-C12 | 1 | 1 |
| IAC-C13 | 1 | 1 |
| IAC-C14 | 1 | 1 |
| IAC-C15 | 1 | 1 |
| IAC-C16 | 1 | 1 |
| IAC873396 | 1 | 1 |
| IAC-C17 | 1 | 1 |
| IAC-C18 | 2 | 2 |
| IAC-C19 | 2 | 2 |
| IAC-C20 | 3 | 3 |
| IAC-C21 | 2 | 2 |
| IAC-C22 | 2 | 2 |
| IAC-C23 | 2 | 2 |
| IAC-C24 | 2 | 2 |
| IAC-C25 | 3 | 3 |
| IAC-C26 | 2 | 2 |
| IAC-C27 | 2 | 2 |
| IAC-C28 | 2 | 2 |
| IAC-C29 | 2 | 2 |
| IAC-C30 | 1 | 1 |
| IAC-C31 | 1 | 2 |
| IAC-C32 | 2 | 2 |
| IAC-C33 | 2 | 2 |
| IAC-C34 | 3 | 3 |
| IAC-C35 | 2 | 2 |
| IAC-C36 | 2 | 2 |
| IAC-C37 | 2 | 2 |
| IAC-C38 | 2 | 2 |
| IAC-C39 | 2 | 2 |
| IAC-C40 | 3 | 3 |
| IAC-C41 | 2 | 2 |
| IAC-C42 | 1 | 1 |
| IAC-C43 | 3 | 3 |
| IAC-C44 | 2 | 2 |
| IAC-C45 | 3 | 3 |
| IAC-C46 | 2 | 2 |
| IACSP013127 | 1 | 1 |
| IAC-C47 | 2 | 2 |
| IACSP015503 | 1 | 1 |
| IAC-C48 | 2 | 2 |
| IAC-C49 | 2 | 2 |
| IAC-C50 | 2 | 2 |
| IAC-C51 | 2 | 2 |
| IAC-C52 | 3 | 3 |
| IAC-C53 | 2 | 2 |
| IAC-C54 | 2 | 2 |
| IAC-C55 | 2 | 2 |
| IAC-C56 | 2 | 2 |
| IAC-C57 | 3 | 3 |
| IAC-C58 | 2 | 2 |
| IAC-C59 | 2 | 2 |
| IAC-C60 | 2 | 2 |
| IAC-C61 | 2 | 2 |
| IAC-C62 | 2 | 2 |
| IAC-C63 | 2 | 2 |
| IAC-C64 | 2 | 2 |
| IAC-C65 | 2 | 2 |
| IAC-C66 | 2 | 2 |
| IAC-C67 | 2 | 2 |
| IAC-C68 | 2 | 2 |
| IAC-C69 | 2 | 2 |
| IAC-C70 | 2 | 2 |
| IAC-C71 | 2 | 2 |
| IAC-C72 | 2 | 2 |
| IAC-C73 | 2 | 2 |
| IAC-C74 | 2 | 2 |
| IAC-C75 | 2 | 2 |
| IAC-C76 | 2 | 2 |
| IAC-C77 | 1 | 1 |
| IAC-C78 | 2 | 2 |
| IACSP932060 | 1 | 1 |
| IACSP933046 | 2 | 2 |
| IAC-C79 | 1 | 1 |
| IACSP942094 | 1 | 1 |
| IACSP942101 | 1 | 1 |
| IACSP944004 | 1 | 1 |
| IAC-C80 | 2 | 2 |
| IAC-C81 | 2 | 2 |
| IACSP955000 | 1 | 1 |
| IACSP955037 | 2 | 2 |
| IACSP955094 | 1 | 1 |
| IAC-C82 | 2 | 2 |
| IAC-C83 | 2 | 2 |
| IACSP962042 | 1 | 1 |
| IAC-C84 | 2 | 2 |
| IAC-C85 | 2 | 2 |
| IAC-C86 | 3 | 2 |
| IAC-C87 | 2 | 2 |
| IAC-C88 | 2 | 2 |
| IAC-C89 | 2 | 2 |
| IAC-C90 | 3 | 3 |
| IAC-C91 | 3 | 3 |
| IAC-C92 | 1 | 1 |
| IAC-C93 | 2 | 2 |
| IAC-C94 | 2 | 2 |
| IAC-C95 | 2 | 2 |
| IAC-C96 | 3 | 3 |
| IAC-C97 | 1 | 1 |
| IAC-C98 | 2 | 2 |
| IAC-C99 | 2 | 2 |
| IAC-C100 | 2 | 2 |
| IAC-C101 | 2 | 2 |
| IAC-C102 | 2 | 2 |
| IAC-C103 | 2 | 2 |
| KASSOER | 1 | 1 |
| KRAKATAU | 2 | 2 |
| NA5679 | 1 | 1 |
| NCO310 | 1 | 1 |
| NG5712 | 2 | 2 |
| NG57213 | 1 | 2 |
| POJ2878 | 1 | 1 |
| R570 | 1 | 1 |
| RAGNAR | 1 | 1 |
| RB855156 | 1 | 1 |
| RB855453 | 1 | 1 |
| RB867515 | 1 | 1 |
| RB92579 | 1 | 1 |
| RB935744 | 2 | 2 |
| SES205A | 1 | 1 |
| SP716163 | 1 | 1 |
| SP701143 | 1 | 1 |
| SP80185 | 1 | 1 |
| SP803280 | 1 | 1 |
| SP813250 | 1 | 1 |
| SP832847 | 2 | 2 |
| SP835073 | 1 | 3 |
| SP901638 | 1 | 1 |
| SP903414 | 1 | 1 |
| SP911049 | 1 | 1 |
| U55714105 | 1 | 1 |
| US571415 | 1 | 1 |
| VERTIX2 | 1 | 1 |
| VERTIX3 | 1 | 1 |
| VERTIX7 | 1 | 1 |
| VERTIX9 | 1 | 1 |

**Table S3. Functional annotation and corresponding R570 Gene IDs of resistance-associated SNPs detected by GWAS.**

| **SNP ID** | **Annotation** | **Chromosome(s) R570** | **Gene ID** |
| --- | --- | --- | --- |
| 3447 | Protein Reduced Wall Acetylation 3 (RWA3) | Chr7os1 | SoffiXsponR570.7os1g002700 |
|  |  | Chr7C | SoffiXsponR570.07Cg009500 |
|  |  | Chr7B | SoffiXsponR570.07Bg008000 |
|  |  | Chr7A | SoffiXsponR570.07Ag008100 |
| 18183 | SCYTHE/BAT3 | Chr8E | SoffiXsponR570.08Eg106300 |
|  |  | Chr8_5A | SoffiXsponR570.8_5Ag171200 |
|  |  | Chr8A | SoffiXsponR570.08Ag173100 |
|  |  | Chr8D | SoffiXsponR570.08Dg154600 |
|  |  | Chr8_10A | SoffiXsponR570.8_10Ag298000 |
|  |  | Chr8C | SoffiXsponR570.08Cg162300 |
| 56 | Protein DNJ-23 related | Chr10os1 | SoffiXsponR570.10os1g044100 |
|  |  | Chr10D | SoffiXsponR570.10Dg079400 |
|  |  | Chr10C | SoffiXsponR570.10Cg083500 |
|  |  | Chr10A | SoffiXsponR570.10Ag104700 |
|  |  | Chr10os2 | SoffiXsponR570.10os2g000600 |
|  |  | Chr10F | SoffiXsponR570.10Fg000500 |
|  |  | Chr10E | SoffiXsponR570.10Eg025200 |
|  |  | Chr10B | SoffiXsponR570.10Bg140700 |
| 1676 | Pentatricopeptide Repeat Domain (PPR3) | Chr4G | SoffiXsponR570.04Gg010700 |
|  |  | Chr4C | SoffiXsponR570.04Cg009900 |
|  |  | Chr4A | SoffiXsponR570.04Ag010700 |
|  |  | Chr4D | SoffiXsponR570.04Dg010400 |
|  |  | Chr4B | SoffiXsponR570.04Bg010300 |
| 4910 | Nitric Oxide Synthase-interacting Protein | Chr1F | SoffiXsponR570.01Fg214000 |
|  |  | Chr1E | SoffiXsponR570.01Eg326600 |
|  |  | Chr1C | SoffiXsponR570.01Cg266100 |
|  |  | Chr1B | SoffiXsponR570.01Bg375000 |
|  |  | Chr1A | SoffiXsponR570.01Ag418400 |
|  |  | Chr1D | SoffiXsponR570.01Dg272900 |
|  | PSBP-like Protein 2 Chloroplastic (PPL2) | Chr1F | SoffiXsponR570.01Fg213900 |
|  |  | Chr1E | SoffiXsponR570.01Eg326500 |
|  |  | Chr1C | SoffiXsponR570.01Cg266000 |
|  |  | Chr1B | SoffiXsponR570.01Bg374900 |
|  |  | Chr1A | SoffiXsponR570.01Ag418300 |
|  |  | Chr1D | SoffiXsponR570.01Dg272800 |
| 4912 / 4913 | Inosine Triphosphate Pyrophosphatase (ITPA) | Chr1D | SoffiXsponR570.01Dg097900 |
|  |  | Chr1A | SoffiXsponR570.01Ag233100 |
|  |  | Chr6B | SoffiXsponR570.06Bg008500 |
|  |  | Chr6F | SoffiXsponR570.06Fg034500 |
|  |  | Chr6C | SoffiXsponR570.06Cg016900 |
|  |  | Chr6E | SoffiXsponR570.06Eg044500 |
|  |  | Chr6_9A | SoffiXsponR570.6_9Ag133000 |
|  |  | Chr1F | SoffiXsponR570.01Fg032600 |
|  |  | Chr1B | SoffiXsponR570.01Bg215300 |
|  |  | Chr6A | SoffiXsponR570.06Ag038300 |
| 18475 | Outer Envelope Protein 80 Chloroplastic (OEP80) | Chr3os5 | SoffiXsponR570.3os5g001000 |
|  |  | Chr3F | SoffiXsponR570.03Fg086300 |
|  |  | Chr3E | SoffiXsponR570.03Eg067900 |
|  |  | Chr3D | SoffiXsponR570.03Dg081500 |
|  |  | Chr3C | SoffiXsponR570.03Cg073300 |
|  |  | Chr3B | SoffiXsponR570.03Bg076700 |
|  |  | Chr3A | SoffiXsponR570.03Ag078200 |
| 20199 | Gamma-irradiation and Mitomycin C Induced 1 | Chr3B | SoffiXsponR570.03Bg104200 |
|  |  | Chr3B | - |
|  |  | Chr3A | SoffiXsponR570.03Ag122300 |
|  |  | Chr3os1 | SoffiXsponR570.3os1g014500 |
|  |  | Chr3G | SoffiXsponR570.03Gg023200 |
|  |  | Chr3G | - |
|  |  | Chr3F | SoffiXsponR570.03Fg134300 |
|  |  | Chr3D | SoffiXsponR570.03Dg128900 |
|  |  | Chr3C | - |
|  |  | Chr3os2 | SoffiXsponR570.3os2g010500 |
| 26739 / 26740 | Phage-T4 UV Endonuclease | Chr3B | SoffiXsponR570.03Bg008100 |
|  | Reverse transcriptases | Chr1D | SoffiXsponR570.01Dg060700 |
| 32075 | Adenine nucleotide alpha hydrolases-domain containing protein kinase | Chr10os1 | SoffiXsponR570.10os1g017800 |
|  |  | Chr10D | SoffiXsponR570.10Dg053700 |
|  |  | Chr10B | SoffiXsponR570.10Bg119300 |
|  |  | Chr7_10A | SoffiXsponR570.7_10Ag074000 |
|  |  | Chr10C | SoffiXsponR570.10Cg058200 |
|  |  | Chr10A | SoffiXsponR570.10Ag076800 |
|  |  | Chr10A | SoffiXsponR570.10Ag071000 |
|  | DNA-directed RNA polymerases | Chr10os1 | SoffiXsponR570.10os1g017700 |
|  |  | Chr10D | SoffiXsponR570.10Dg053800 |
|  |  | Chr10B | SoffiXsponR570.10Bg119400 |
|  |  | Chr7_10A | SoffiXsponR570.7_10Ag073900 |
|  |  | Chr10C | SoffiXsponR570.10Cg058100 |
|  |  | Chr10A | SoffiXsponR570.10Ag076700 |
|  |  | Chr10A | SoffiXsponR570.10Ag070900 |
| 9291 | E3 ubiquitin-protein ligase (UBR3) | Chr2B | SoffiXsponR570.02Bg069300 |
|  |  | Chr2A | SoffiXsponR570.02Ag071600 |
|  |  | Chr2F | SoffiXsponR570.02Fg071600 |
|  |  | Chr2E | SoffiXsponR570.02Eg048300 |
|  |  | Chr2C | SoffiXsponR570.02Cg070900 |
|  |  | Chr2D | SoffiXsponR570.02Dg068300 |
|  | Lectin Receptor-like Kinase (LecRLK) | Chr2B | - |
|  |  | Chr2A | SoffiXsponR570.02Ag071500 |
|  |  | Chr2F | SoffiXsponR570.02Fg071500 |
|  |  | Chr2E | SoffiXsponR570.02Eg048200 |
|  |  | Chr2C | SoffiXsponR570.02Cg070800 |
|  |  | Chr2D | - |
|  | Leucine-rich Repeat-containing Protein (LRR) | Chr2D | SoffiXsponR570.02Dg068200 |

**Table S4. Predicted coding effects and genomic context of resistance-associated SNPs located within genic regions.** Reference and alternative nucleotide triplets and their corresponding amino acids are presented for coding-region SNPs to indicate synonymous and non-synonymous substitutions, while SNPs located in untranslated or intronic regions are annotated according to their respective genomic context.

| **SNP ID** | **Gene** | **Codon** | | **Amino acid** | |
| --- | --- | --- | --- | --- | --- |
|  |  | **Reference** | **Alternative** | **Reference** | **Alternative** |
| 3447 | Protein Reduced Wall Acetylation 3 (RWA3) | AC**G** | AC**A** | Threonine | Threonine |
| 18183 | SCYTHE/BAT3 | 3'UTR | | | |
| 56 | Protein DNJ-23 related | CT**G** | CT**T** | Leucine | Leucine |
| 1676 | Pentatricopeptide Repeat Domain (PPR3) | G**T**C | G**C**C | Valine | Alanine |
| 4910 | Nitric Oxide Synthase-interacting Protein | 3'UTR | | | |
|  | PSBP-like Protein 2 Chloroplastic (PPL2) | Intron | | | |
| 4912 / 4913 | Inosine Triphosphate Pyrophosphatase (ITPA) | Intron | | | |
| 18475 | Outer Envelope Protein 80 Chloroplastic (OEP80) | Intron | | | |
| 20199 | Gamma-irradiation and Mitomycin C Induced 1 | C**A**T | C**G**T | Histidine | Arginine |
| 26739 / 26740 | Phage-T4 UV Endonuclease | C**TC** | C**GG** | Leucine | Arginine |
|  | Reverse transcriptases |  |  |  |  |

**Table S5. Summary of the transcriptomic datasets used in the analyses.** For each study, the sugarcane genotype and smut resistance phenotype, treatment and fungal isolate, average number of reads before and after trimming, and mean mapping rate obtained with Salmon, calculated from three biological replicates per treatment, are presented.

| **Study** | **Genotype** | **Phenotype** | **Treatment** | **Fungal isolate** | **Mean number of reads** | | **Mean mapping rate (%)** |
| --- | --- | --- | --- | --- | --- | --- | --- |
|  |  |  |  |  | **Before trimming** | **After trimming** |  |
| Schaker *et al*. (2016) | RB925345 | Intermediately  resistant | 5 DAI (Control) | - | 9,250,753 | 8,492,836 | 60.88 |
|  |  |  | 200 DAI (Control) | - | 9,375,709 | 8,208,341 | 67.50 |
|  |  |  | 5 DAI | SSC39 | 9,393,251 | 8,539,963 | 60.28 |
|  |  |  | 200 DAI | SSC39 | 9,502,495 | 8,366,948 | 52.52 |
| McNeil *et al*. (2018) | CP74-2005 | Susceptible | 48 HAI (Control) | - | 19,945,242 | 19,189,646 | 59.92 |
|  |  |  | 48 HAI | NA | 24,921,748 | 22,657,825 | 59.67 |
| Agisha *et al*. (2022) | Co 97009 | Susceptible | 2 DAI (Control) | - | 22,824,341 | 18,606,765 | 19.59 |
|  |  |  | 5 DAI (Control) | - | 18,346,618 | 16,227,950 | 31.77 |
|  |  |  | 60 DAI (Control) | - | 29,080,106 | 24,766,767 | 31.33 |
|  |  |  | 2 DAI | Ss97009 | 27,297,424 | 23,016,317 | 16.49 |
|  |  |  | 5 DAI | Ss97009 | 24,753,823 | 21,248,022 | 27.26 |
|  |  |  | 60 DAI | Ss97009 | 21,303,014 | 18,121,446 | 25.36 |
|  |  |  | 2 DAI | SsV89101 | 29,260,195 | 24,774,124 | 25.17 |
|  |  |  | 5 DAI | SsV89101 | 20,284,453 | 17,425,854 | 31.16 |
|  |  |  | 60 DAI | SsV89101 | 24,772,995 | 20,877,323 | 29.68 |
| Wu *et al.*  (2022) | YT93-159 | Resistant | 0 DAI (Control) | - | 38,414,164 | 35,885,717 | 67.58 |
|  |  |  | 1 DAI | NA | 36,379,277 | 36,913,937 | 68.21 |
|  |  |  | 2 DAI | NA | 35,648,961 | 33,647,560 | 67.41 |
|  |  |  | 3 DAI | NA | 39,388,185 | 36,718,214 | 66.94 |
|  |  |  | 4 DAI | NA | 41,678,923 | 37,872,358 | 67.32 |
|  |  |  | 5 DAI | NA | 36,324,866 | 34,023,822 | 66.44 |
|  | ROC22 | Susceptible | 0 DAI (Control) | - | 40,984,745 | 33,542,000 | 66.34 |
|  |  |  | 1 DAI | NA | 33,694,651 | 32,582,964 | 68.18 |
|  |  |  | 2 DAI | NA | 35,103,407 | 32,838,462 | 67.81 |
|  |  |  | 3 DAI | NA | 50,408,450 | 38,713,468 | 68.36 |
|  |  |  | 4 DAI | NA | 40,574,575 | 31,485,537 | 66.94 |
|  |  |  | 5 DAI | NA | 34,705,187 | 32,579,724 | 68.95 |

DAI: Days after inoculation. HAI: Hours after inoculation.

**Table S7. Results of the differential gene expression analyses performed using the previously generated and processed transcriptomic data.** Comparisons between control and infected samples at different time points post-infection are presented, indicating the total number of differentially expressed genes (DEGs), also distinguished between up-and downregulated genes.

| **Study** | **Comparison** | **DEGs** | **Upregulated** | **Downregulated** |
| --- | --- | --- | --- | --- |
| Schaker *et al*. (2016) | 5 DAI vs. Control 5 DAI | 76 | 40 | 36 |
|  | 200 DAI vs. Control 200 DAI | 1,720 | 1,005 | 715 |
| McNeil *et al.* (2018) | 48 HPI vs. Control 48HPI | 115 | 77 | 38 |
| Agisha *et al.* (2022) | Ss97009 2 DAI vs. Control 2 DAI | 1,187 | 754 | 433 |
|  | Ss97009 5 DAI vs. Control 5 DAI | 388 | 315 | 73 |
|  | Ss97009 60 DAI vs. Control 60 DAI | 1,395 | 1,116 | 279 |
|  | SsV89101 2 DAI vs. Control 2 DAI | 1,231 | 785 | 446 |
|  | SsV89101 5 DAI vs. Control 5 DAI | 2,224 | 1,081 | 1,143 |
|  | SsV89101 60 DAI vs. Control 60 DAI | 509 | 355 | 154 |
| Wu *et al.* (2022) | ROC22 1 DAI vs. Control 0 DAI | 2,634 | 2,315 | 319 |
|  | ROC22 2 DAI vs. Control 0 DAI | 4,535 | 3,453 | 1,082 |
|  | ROC22 3 DAI vs. Control 0 DAI | 2,316 | 1,846 | 470 |
|  | ROC22 4 DAI vs. Control 0 DAI | 4,018 | 3,157 | 861 |
|  | ROC22 5 DAI vs. Control 0 DAI | 4,433 | 3,490 | 943 |
|  | YT93 1 DAI vs. Control 0 DAI | 1,882 | 1,654 | 228 |
|  | YT93 2 DAI vs. Control 0 DAI | 1,842 | 1,720 | 122 |
|  | YT93 3 DAI vs. Control 0 DAI | 4,871 | 4,032 | 839 |
|  | YT93 4 DAI vs. Control 0 DAI | 8,814 | 7,755 | 1,059 |
|  | YT93 5 DAI vs. Control 0 DAI | 11,234 | 8,224 | 3,010 |
| Wu *et al.* (2022) | YT93 1 DAI vs. ROC22 1 DAI | 35,539 | 18,670 | 16,869 |
|  | YT93 2 DAI vs. ROC22 2 DAI | 28,231 | 15,792 | 12,439 |
|  | YT93 3 DAI vs. ROC22 3 DAI | 26,114 | 15,281 | 10,833 |
|  | YT93 4 DAI vs. ROC22 4 DAI | 32,858 | 18,559 | 14,299 |
|  | YT93 5 DAI vs. ROC22 5 DAI | 30,670 | 17,101 | 13,569 |

DAI: Days after inoculation. HAI: Hours after inoculation.

**Table S8. Overrepresented coexpression network modules for each phenotype group.** Expected and observed numbers of differentially expressed genes, together with nominal and Bonferroni-adjusted *p* values, are presented.

| **Phenotype** | **Module** | **Expected** | **Observed** | ***p*Value** | **Adjusted *p* value** |
| --- | --- | --- | --- | --- | --- |
| Resistant | green | 748.98 | 1076 | 1.83e-35 | 6.24e-34 |
|  | yellow | 714.31 | 1285 | 3.47e-101 | 1.18e-99 |
|  | steelblue | 62.26 | 91 | 0.00018 | 0.0063 |
|  | saddlebrown | 94.92 | 221 | 1.83e-32 | 6.23e-31 |
|  | greenyellow | 256.84 | 460 | 7.81e-35 | 2.65e-33 |
|  | lightyellow | 95.89 | 439 | 1.54e-175 | 5.23e-174 |
|  | white | 38.35 | 130 | 2.77e-36 | 9.41e-35 |
|  | skyblue | 83.72 | 404 | 2.13e-171 | 7.24e-170 |
|  | grey | 89.85 | 134 | 2.49e-06 | 8.46e-05 |
| Intermediately  resistant | red | 77.54 | 114 | 3.30e-05 | 0.00112 |
|  | salmon | 25.43 | 50 | 7.38e-06 | 0.00025 |
|  | greenyellow | 38.71 | 169 | 7.98e-58 | 2.71e-56 |
|  | skyblue | 12.62 | 69 | 1.25e-29 | 4.27e-28 |
| Susceptible | salmon | 143.3 | 189 | 7.05e-05 | 0.0024 |
|  | steelblue | 52.87 | 95 | 2.62e-08 | 8.91e-07 |
|  | black | 406.98 | 626 | 1.24e-27 | 4.21e-26 |
|  | blue | 831.81 | 967 | 2.43e-07 | 8.26e-06 |
|  | cyan | 145.67 | 223 | 2.02e-10 | 6.87e-09 |
|  | midnightblue | 124.63 | 180 | 5.53e-07 | 1.88e-05 |
|  | royalblue | 57.78 | 128 | 2.52e-17 | 8.58e-16 |
|  | lightcyan | 72.72 | 177 | 3.85e-28 | 1.31e-26 |
|  | darkolivegreen | 11.53 | 45 | 9.55e-16 | 3.25e-14 |
|  | violet | 11.6 | 44 | 8.13e-15 | 2.76e-13 |

**Table S9. Differentially expressed hub genes and their corresponding module membership (MM) values.** Entries marked with “-” indicate genes that showed temporal differential expression within the same genotype but no significant differential expression between the resistant genotype YT93 and the susceptible genotype ROC22, and therefore were not classified as either up- or downregulated.

| **Module** | **Annotation** | **MM** | **R *vs* S** |
| --- | --- | --- | --- |
| black | CCHC-type zinc finger and BAR domain-containing protein | 0.976 | Up |
| darkolivegreen | Glycine-rich protein | 0.974 | Up |
| darkred | Large subunit ribosomal protein L7Ae (RP-L7Ae) | 0.972 | Up |
| lightyellow | Cysteine-rich receptor-like protein kinase 25 | 0.861 | Up |
| red | Clathrin heavy chain (CLTC) | 0.870 | Up |
| skyblue | HVA22-like protein E | 0.961 | Up |
| tan | Cytosol alanyl aminopeptidase / soluble alanyl aminopeptidase. | 0.945 | Up |
| turquoise | P-LOOP containing nucleoside triphosphate hydrolases superfamily | 0.943 | Up |
| violet | Unannotated | 0.921 | Up |
| white | Sulfate transporter 3.1-related | 0.945 | Up |
| yellow | Chaperone protein DNAJ A6, Chloroplastic | 0.946 | Up |
| blue | Phytocyanin domain-containing protein | 0.971 | Down |
| cyan | 26S Proteasome regulatory subunit T3 (PSMC4, RPT3) | 0.915 | Down |
| darkgreen | WD40 repeat-like superfamily protein | 0.942 | Down |
| darkgrey | Trafficking protein particle complex subunit 10 (TRAPPC10) | 0.932 | Down |
| darkorange | Protein LIN-9 Homolog | 0.951 | Down |
| darkturquoise | Protein MEI2-like 4 | 0.889 | Down |
| lightgreen | Transmembrane superfamily member 8-related | 0.943 | Down |
| magenta | Serine Threonine-protein phosphatase PP2A | 0.976 | Down |
| paleturquoise | Oxysterol-binding Protein-related Protein 3C | 0.922 | Down |
| purple | Eukaryotic initiation factor 4A-III | 0.935 | Down |
| steelblue | 3-ketoacyl-COA synthase 5-related | 0.955 | Down |
| greenyellow | Flavonoid 3'-monooxygenase | 0.980 | - |
| royalblue | Glutathione S-transferase F13 | 0.962 | - |
| lightcyan | Bulb-type lectin receptor-like kinase / PAN domain | 0.958 | - |
| saddlebrown | ABC Transporter domain-containing protein | 0.947 | - |

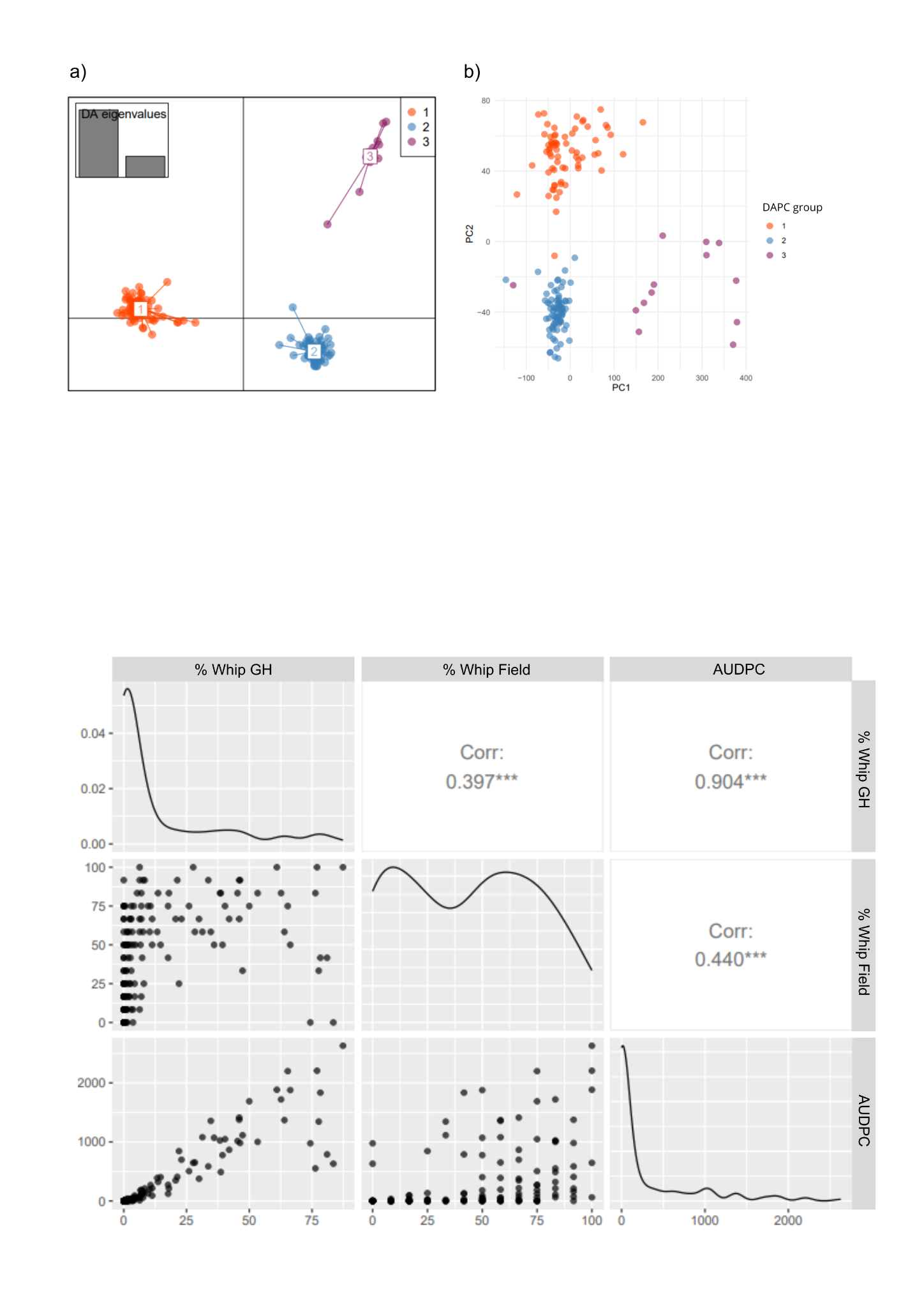

**Fig. S1.** **Distribution and correlation plots among the smut resistance phenotypic traits analysed: percentage of whip emission under greenhouse conditions (% Whip GH), percentage of whip emission under field conditions (% Whip Field), and Area Under the Disease Progress Curve (AUDPC).** The diagonal displays the frequency distributions of each variable; the lower panel shows pairwise scatter plots; and the upper panel presents Pearson correlation coefficients (R) with significance level *p* < 0.001 (***).

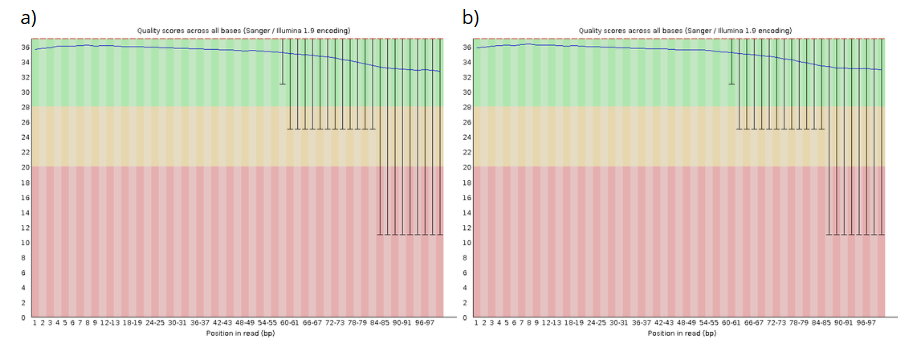

**Fig. S2.** **Plot of the mean base quality scores along the reads for lanes 1 (a) and 2 (b).**

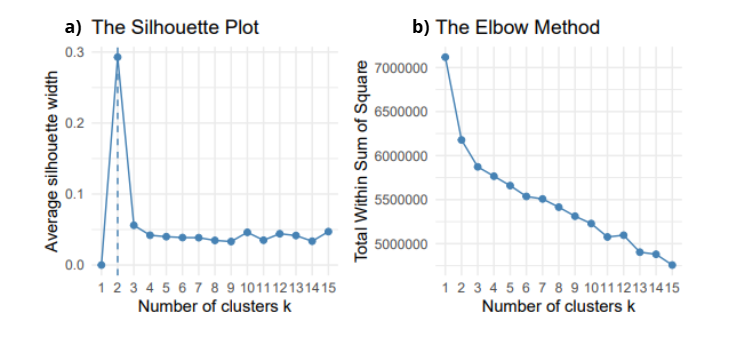

**Fig. S3.** **Determination of the optimal number of clusters (k) in the panel obtained from Principal Component Analysis (PCA) followed by K-means clustering: (a) silhouette method and (b) elbow method.**

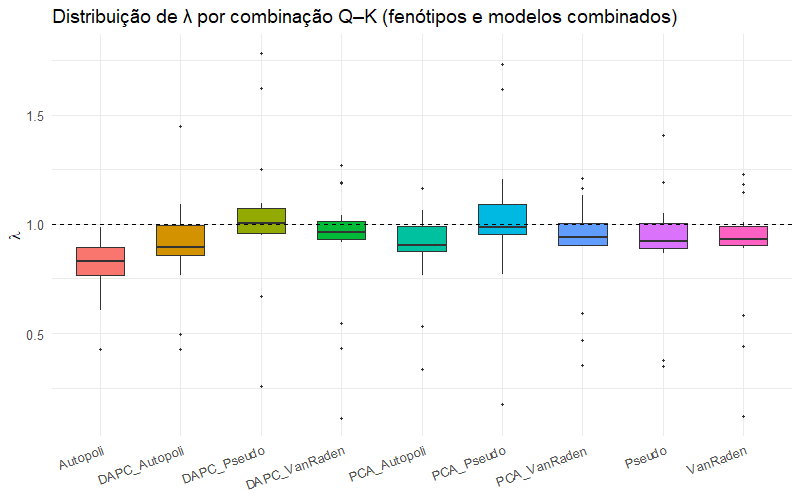

**Fig. S4.** **Distribution of λ values for the different combinations of genetic structure (Q) and kinship (K) matrices tested in the genome-wide association analyses.**

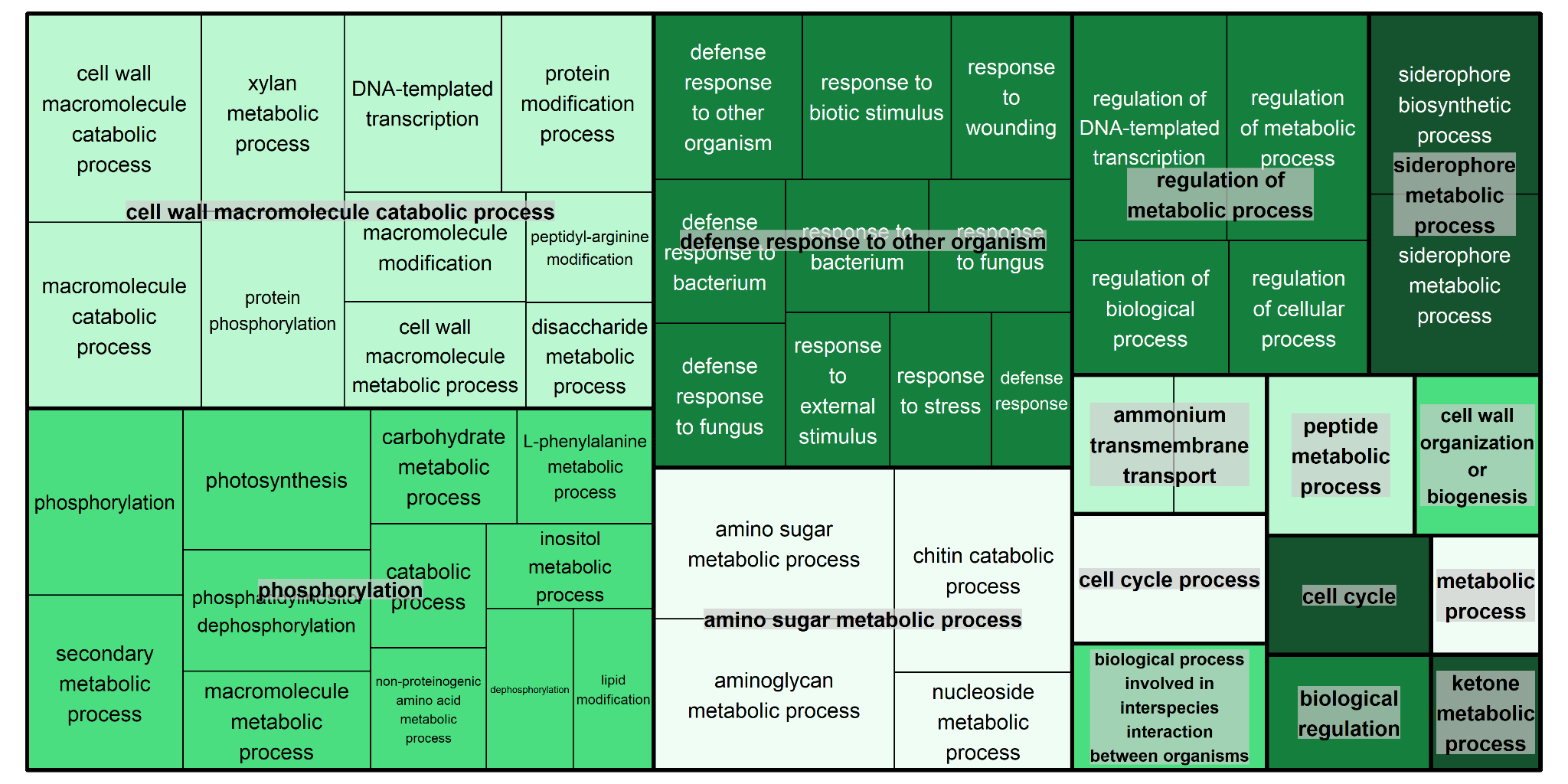

**Fig. S5. Functional categorization of significantly enriched Gene Ontology (GO) biological process terms associated with the complete DEG set of smut-resistant sugarcane genotypes, summarized using ReviGO.**

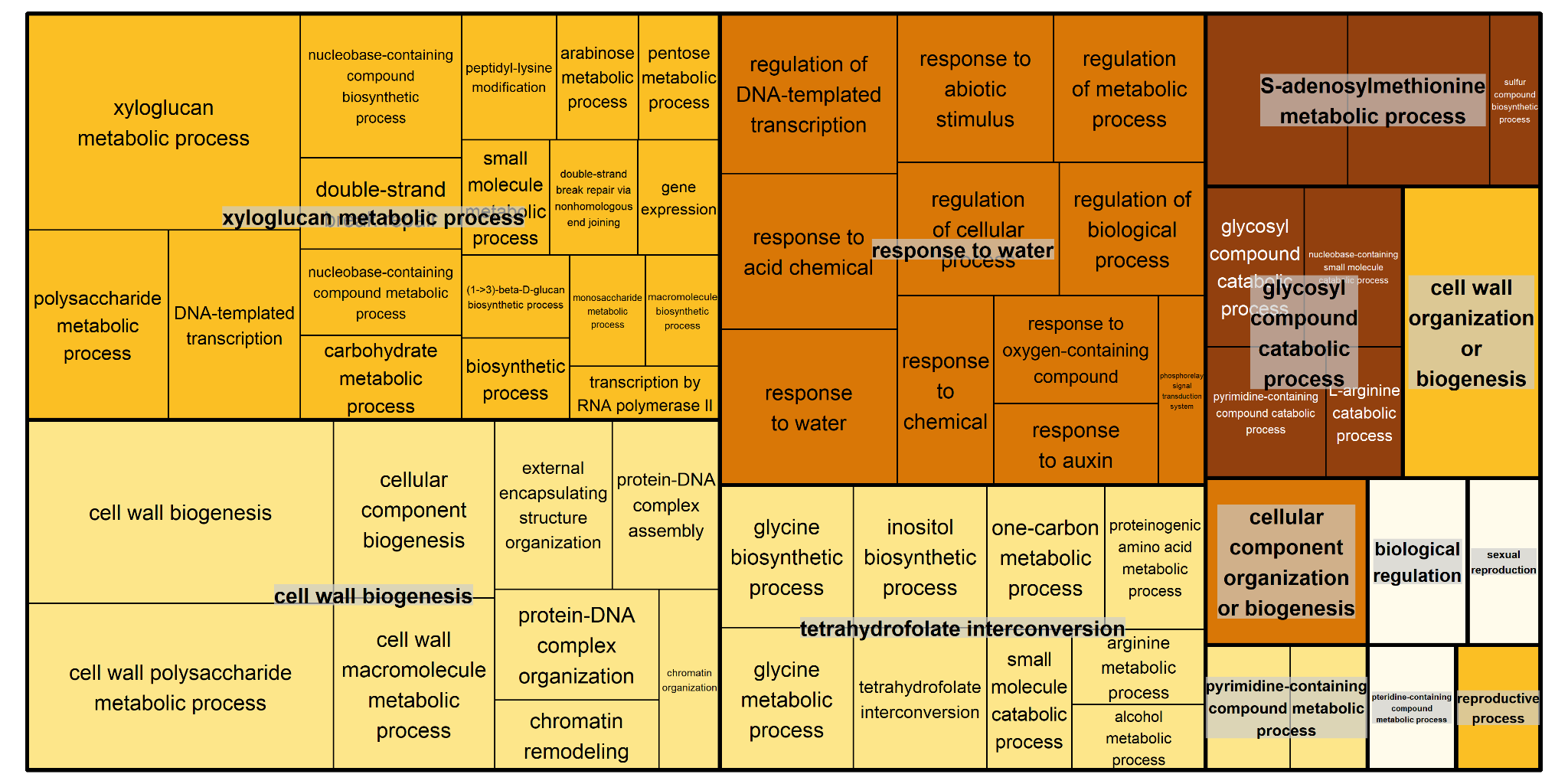

**Fig. S6. Functional categorization of significantly enriched Gene Ontology (GO) biological process terms associated with the complete DEG set of intermediately smut-resistant sugarcane genotypes, summarized using ReviGO.**

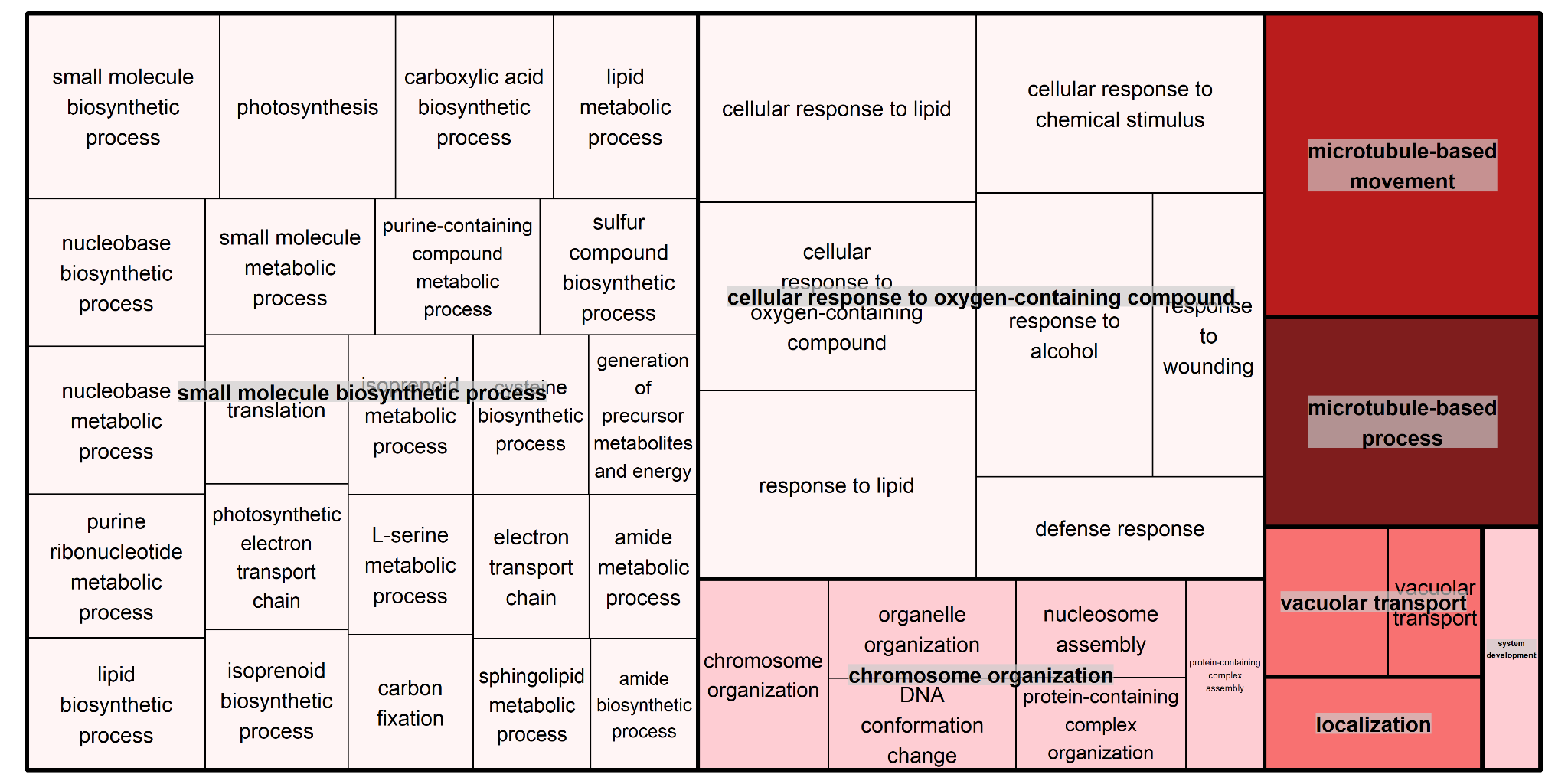

**Fig. S7. Functional categorization of significantly enriched Gene Ontology (GO) biological process terms associated with the complete DEG set of smut-susceptible sugarcane genotypes, summarized using ReviGO.**

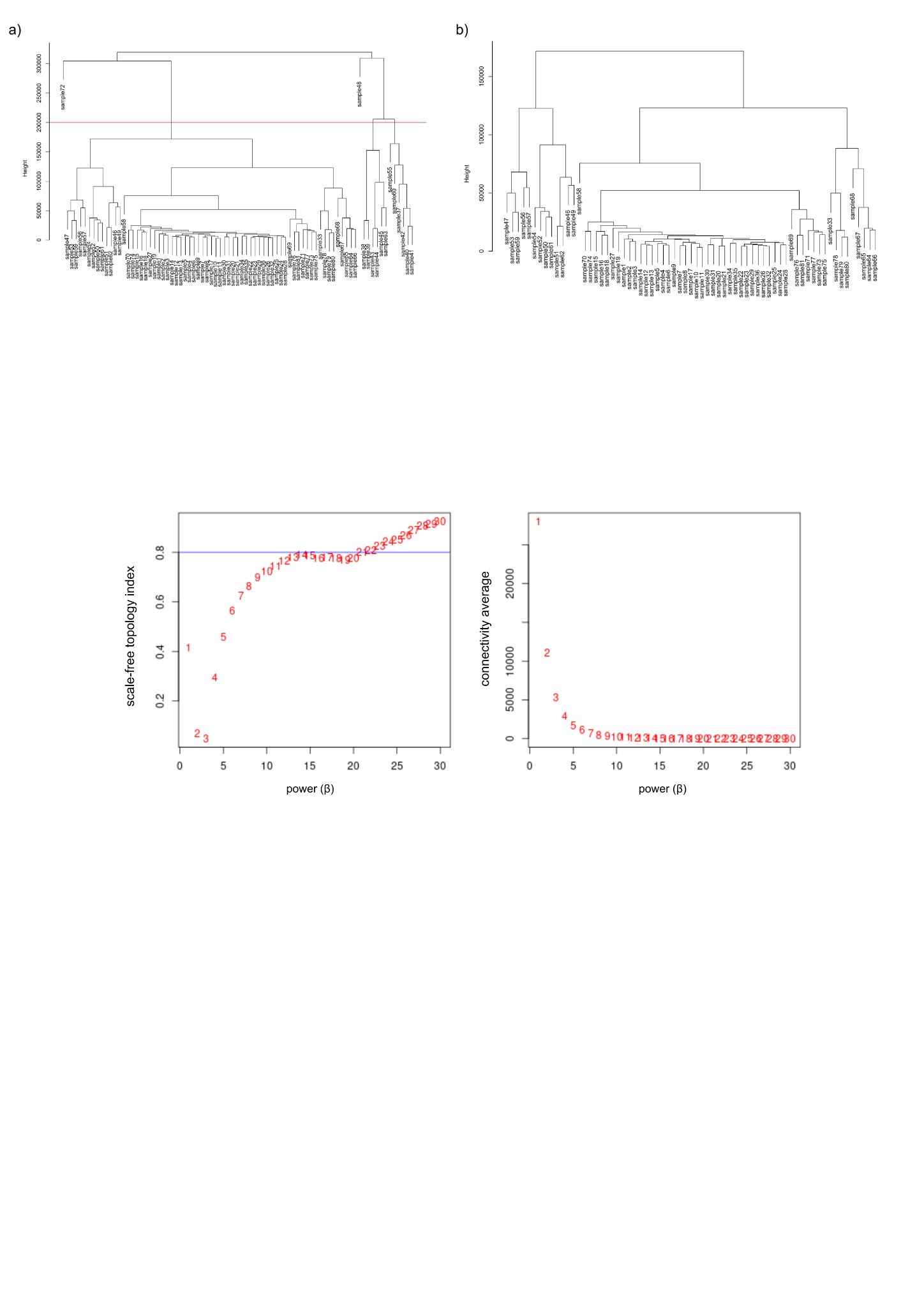

**Fig. S8. Sample clustering for outlier detection for weighted gene coexpression network analyses.** (a) Hierarchical clustering of all 81 samples before filtering; (b) Hierarchical clustering after outlier removal.

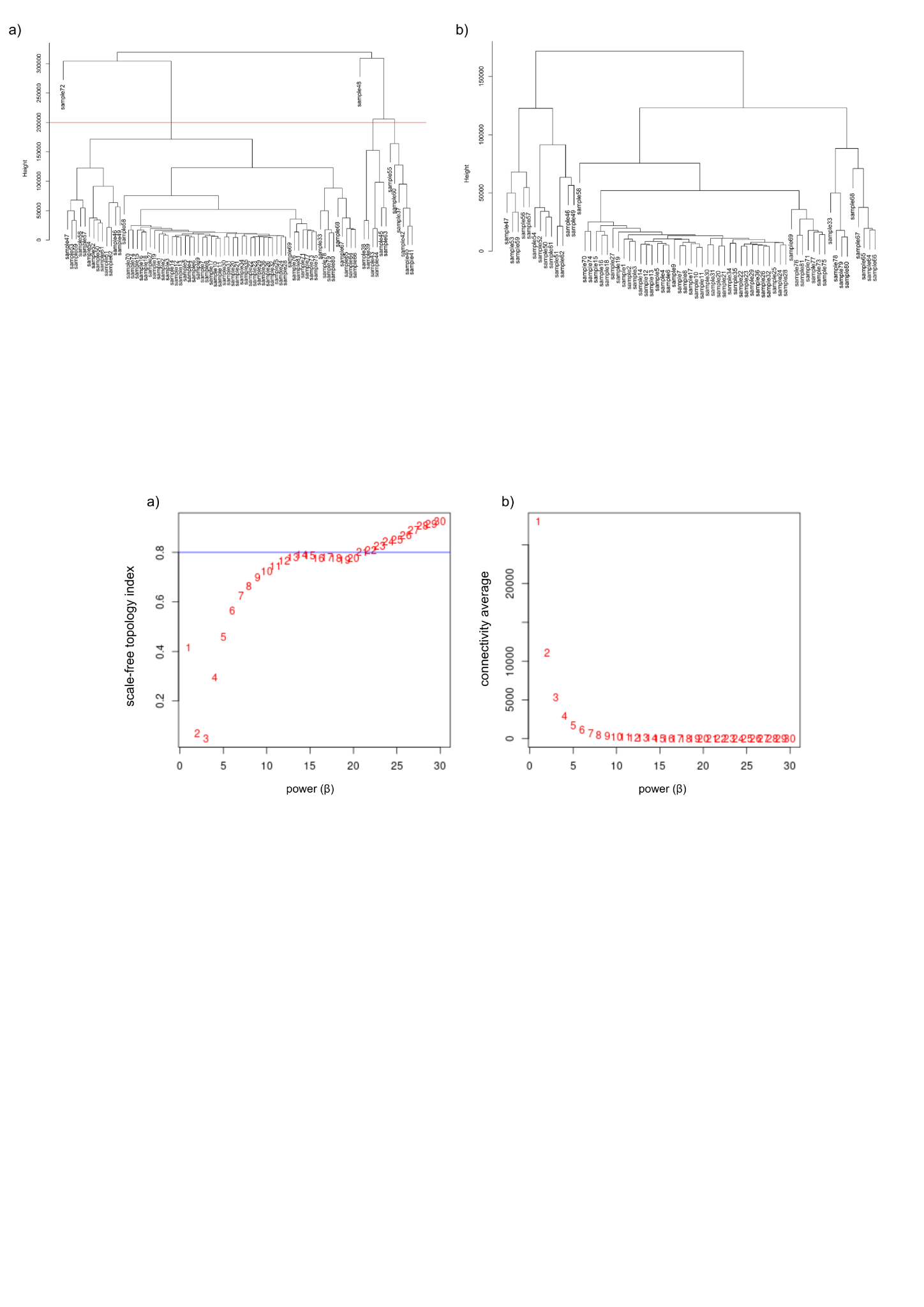

**Fig. S9. Soft-thresholding power analysis for weighted gene coexpression network construction.** (a) Scale-free topology fit index (β) and (b) mean connectivity across different soft-thresholding powers.

**Fig. S10. REVIGO TreeMaps summarizing Gene Ontology functional enrichment of co-expression modules showing overrepresentation of differentially expressed genes exclusive to smut-resistant sugarcane genotypes.** Panels correspond to the following modules: (a) green, (b) yellow, (c) steelblue, (d) saddlebrown, (e) greenyellow, (f) lightyellow, (g) white, (h) skyblue, and (i) grey. Rectangle size is proportional to the associated enrichment value.

**a)**

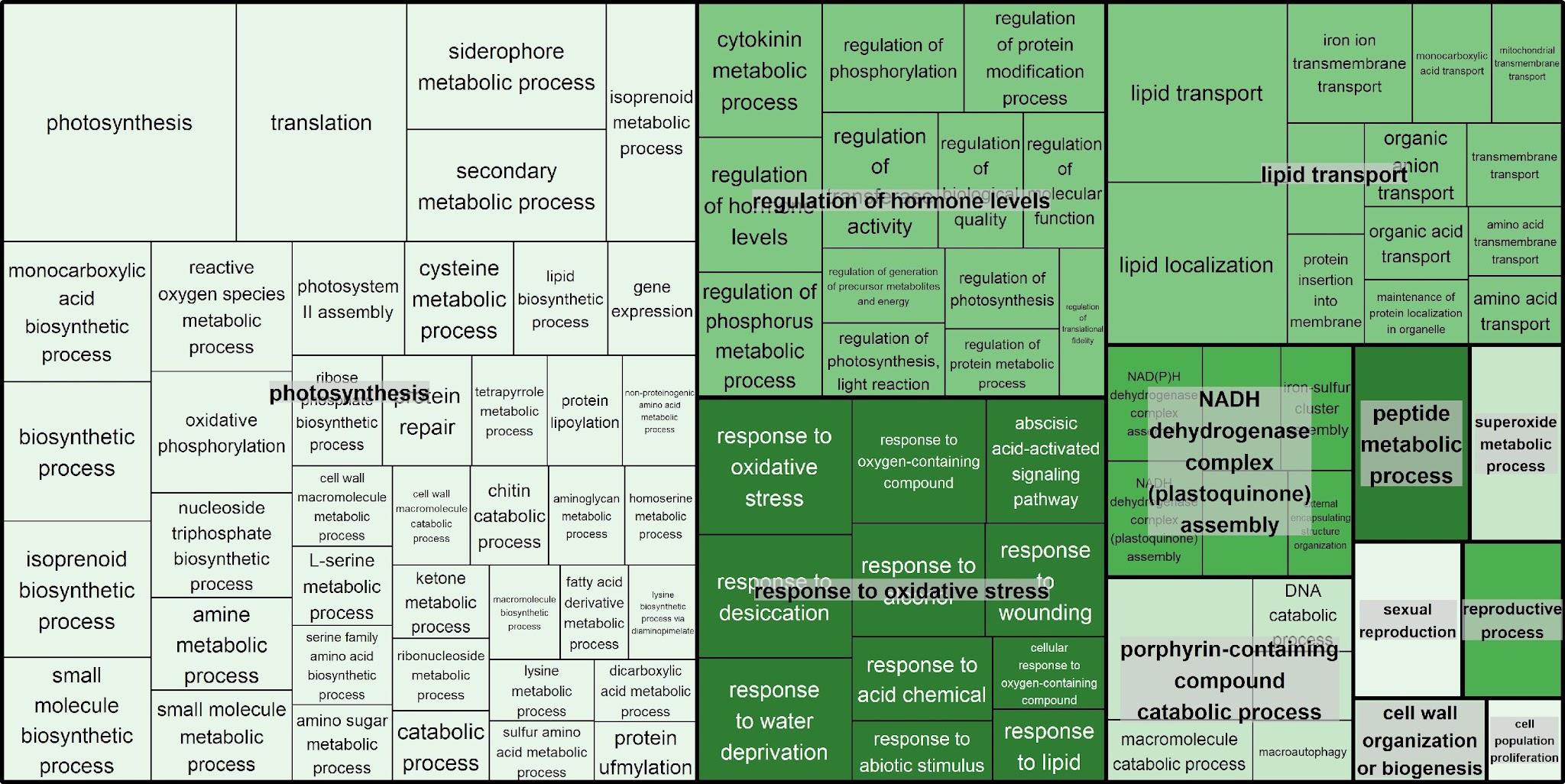

**b)**

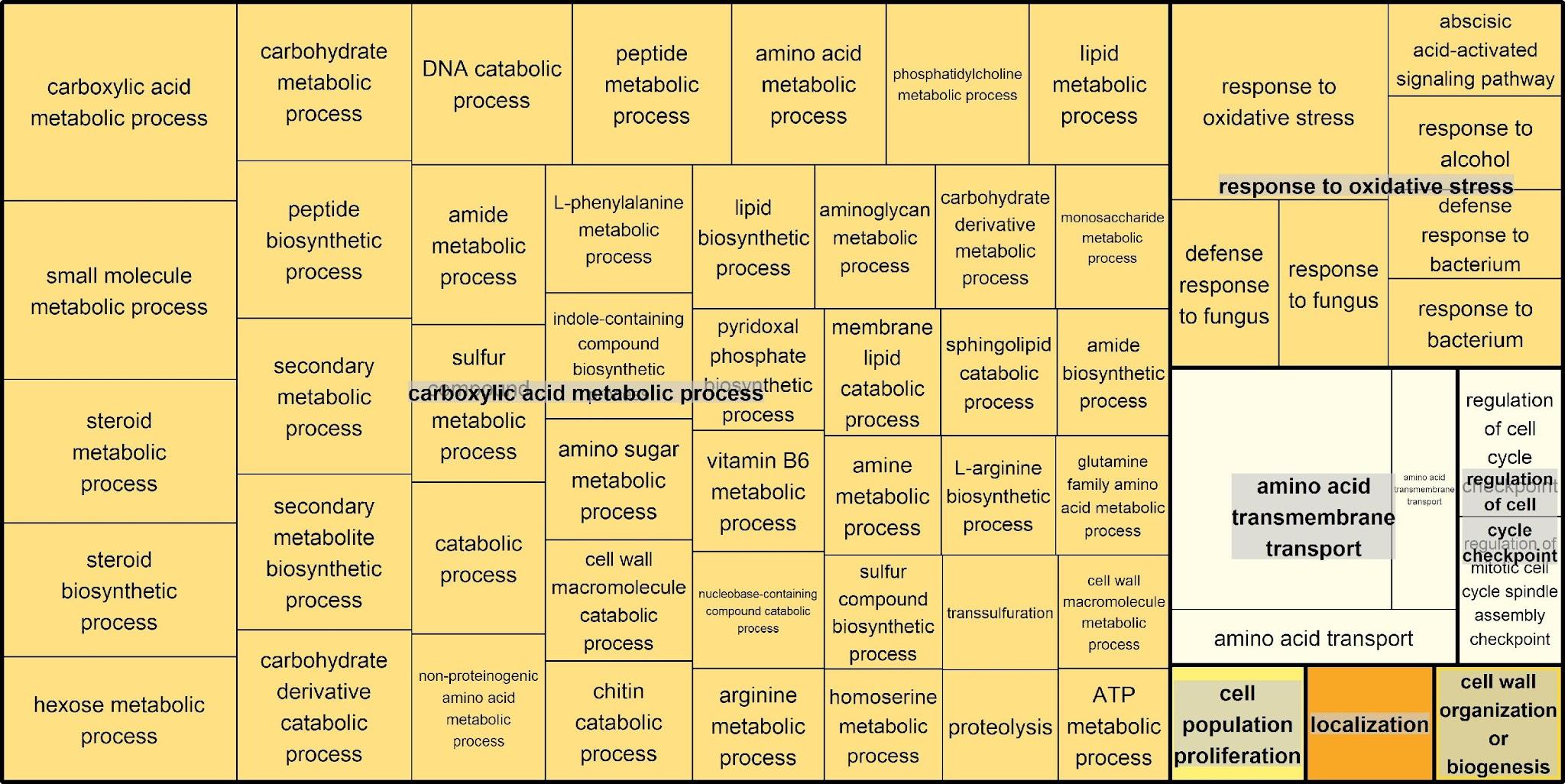

**c)**

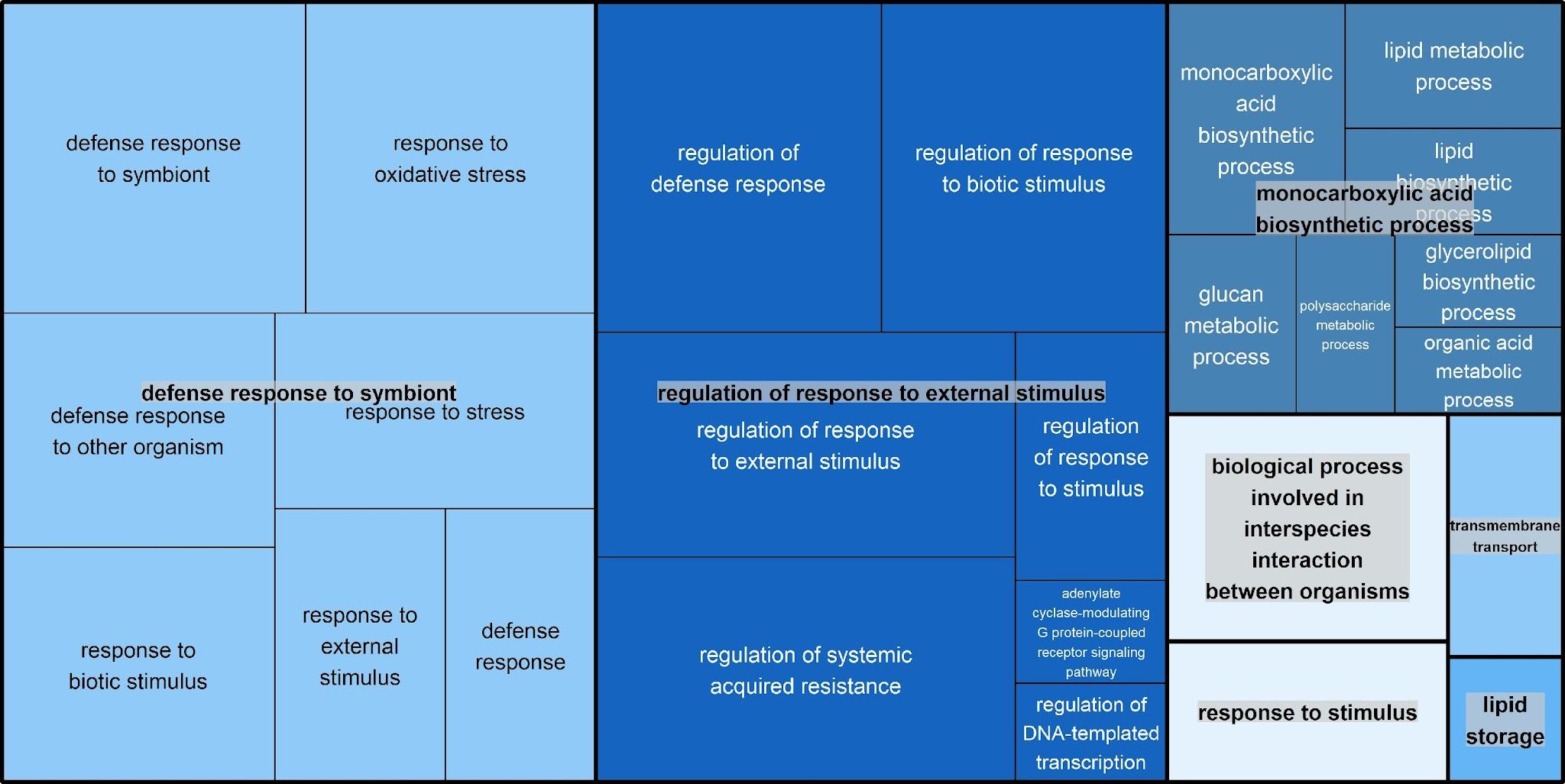

**d)**

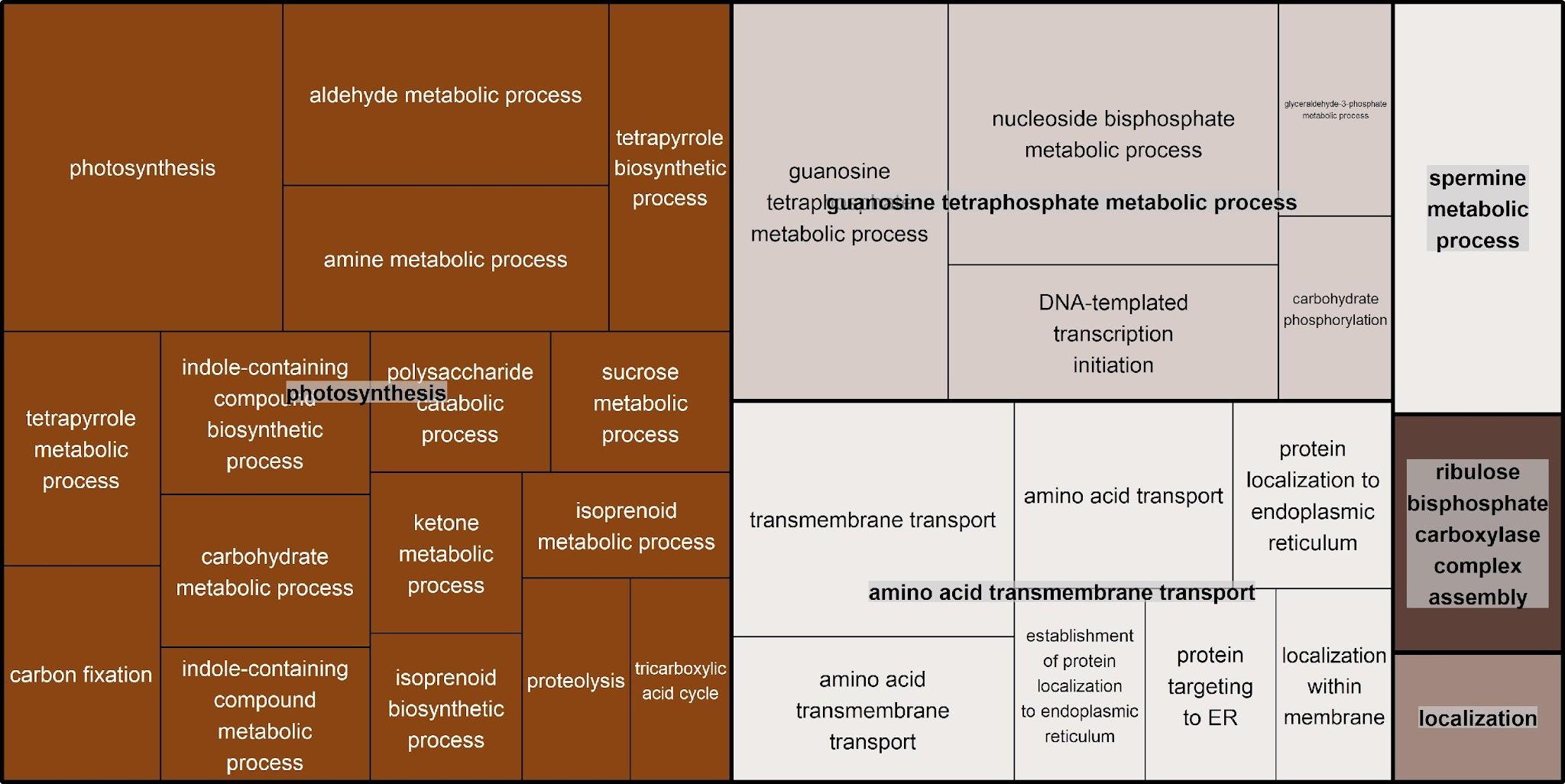

**e)**

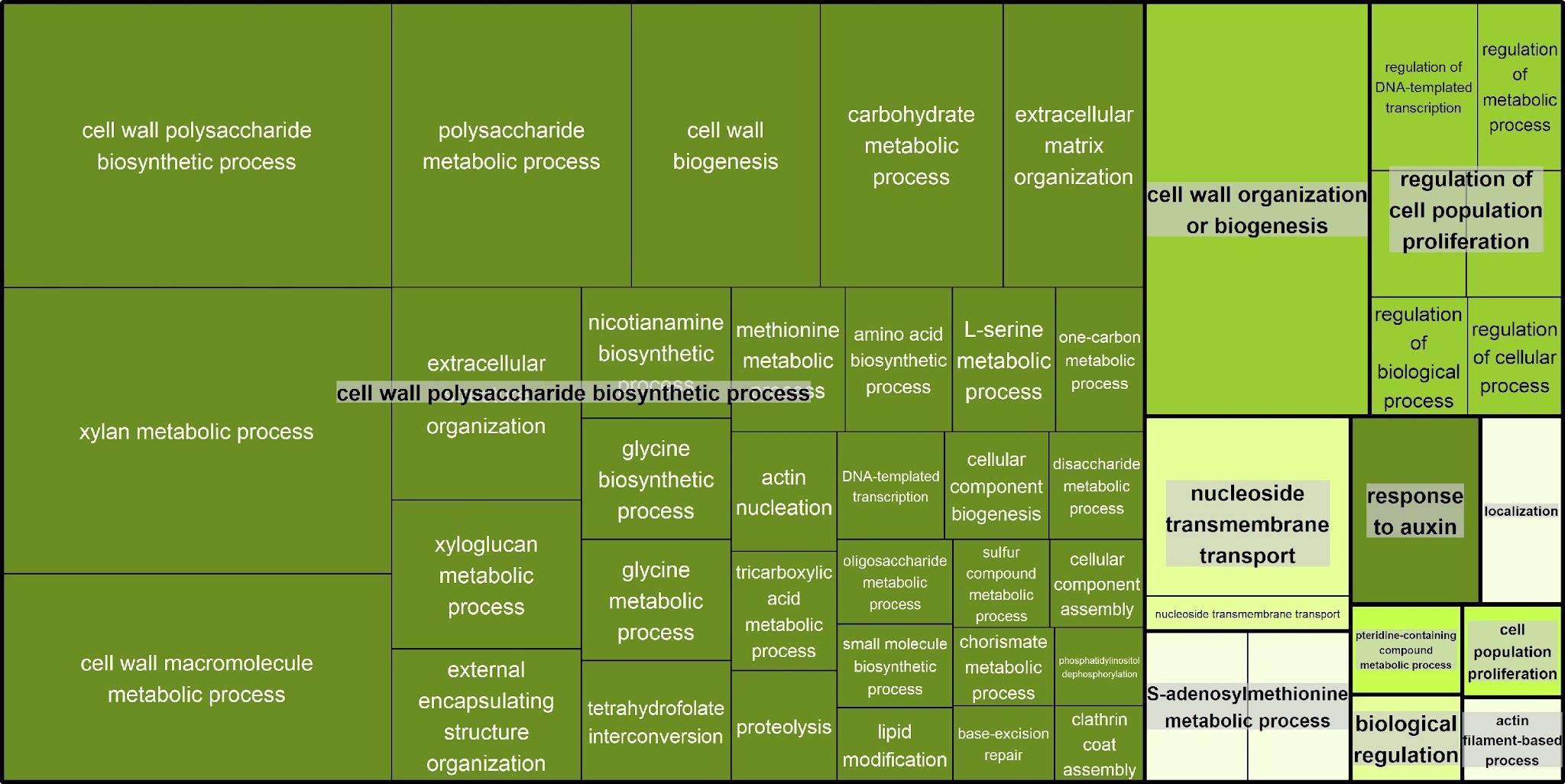

**f)**

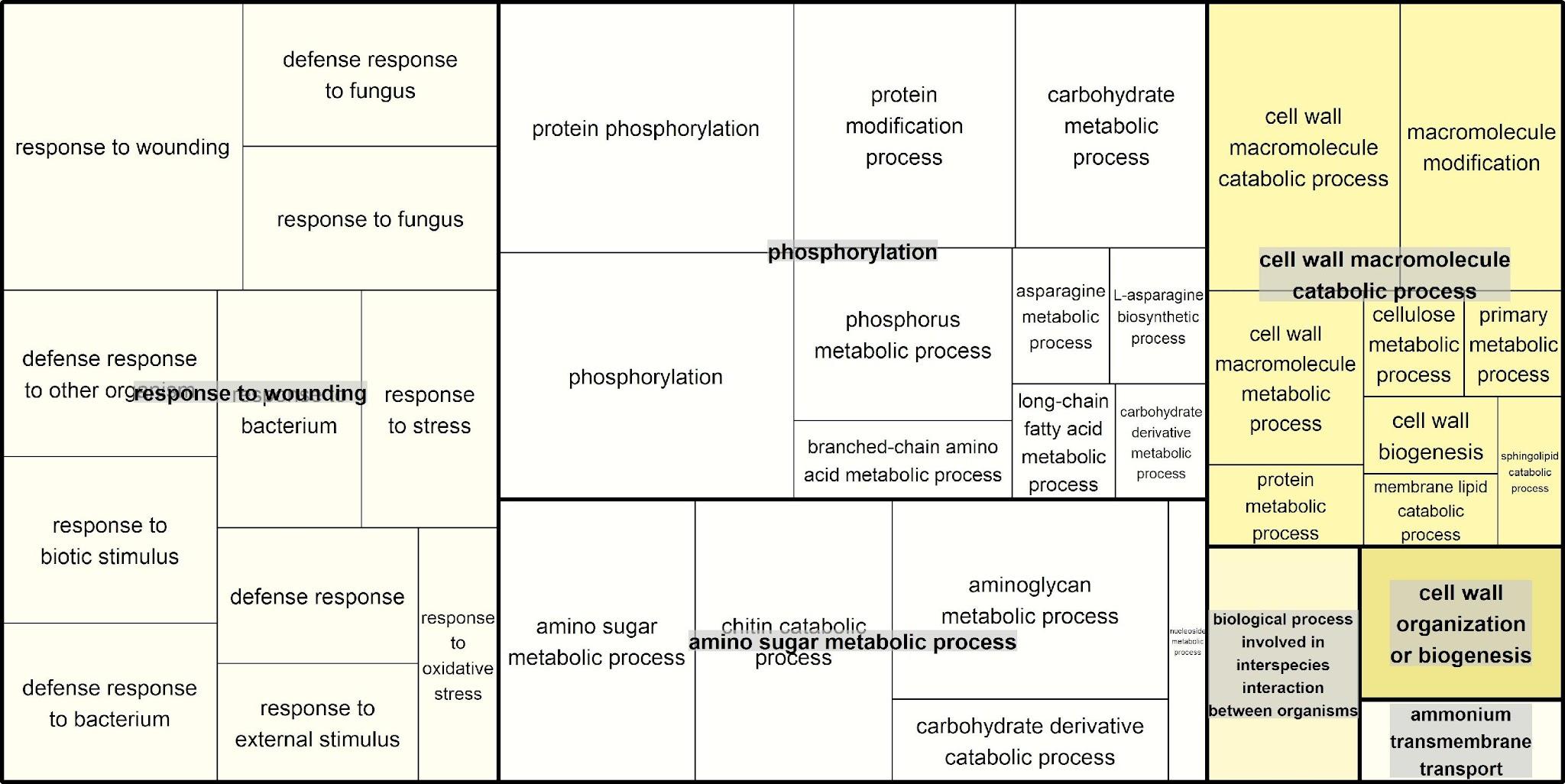

**g)**

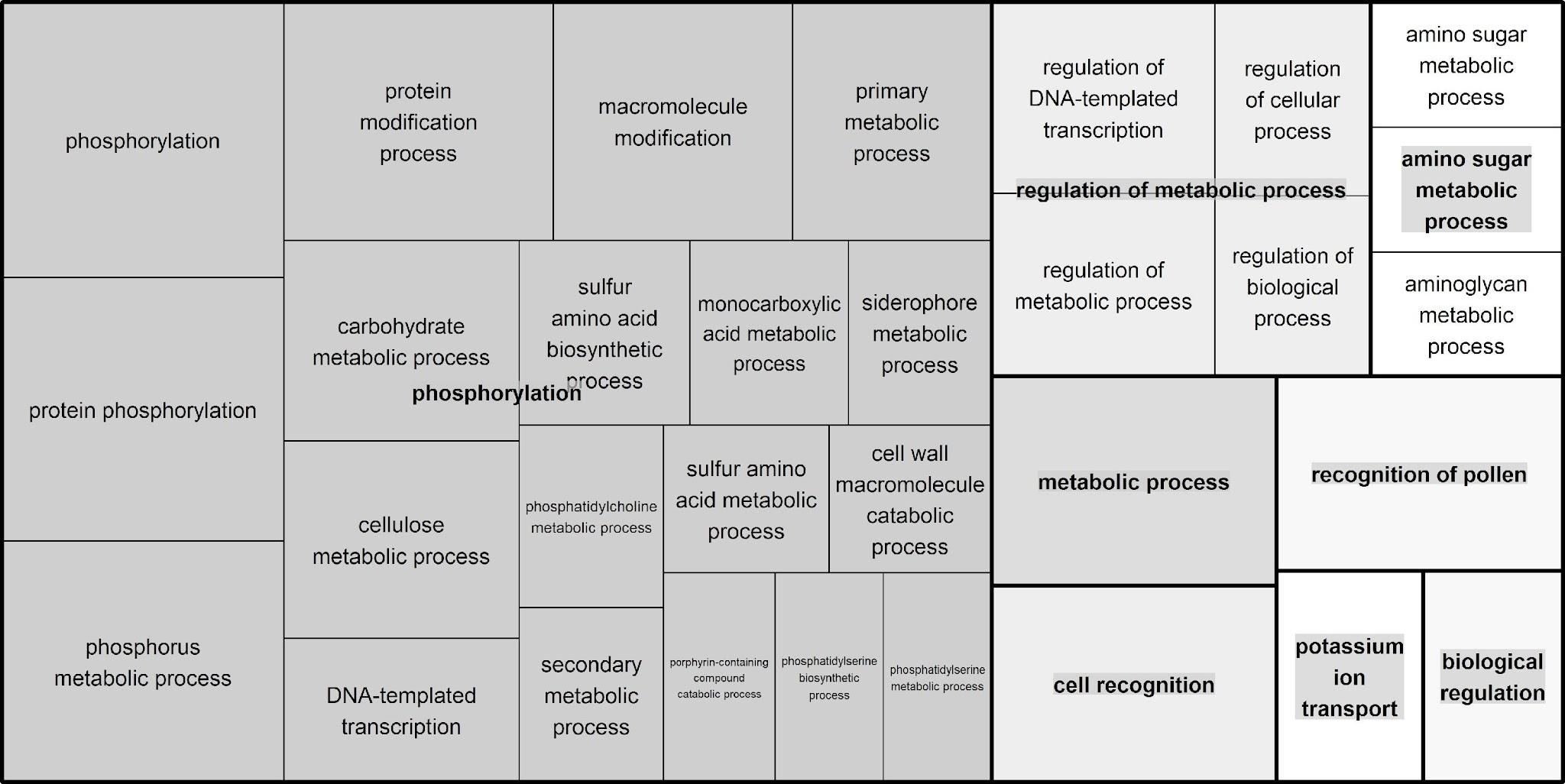

**h)**

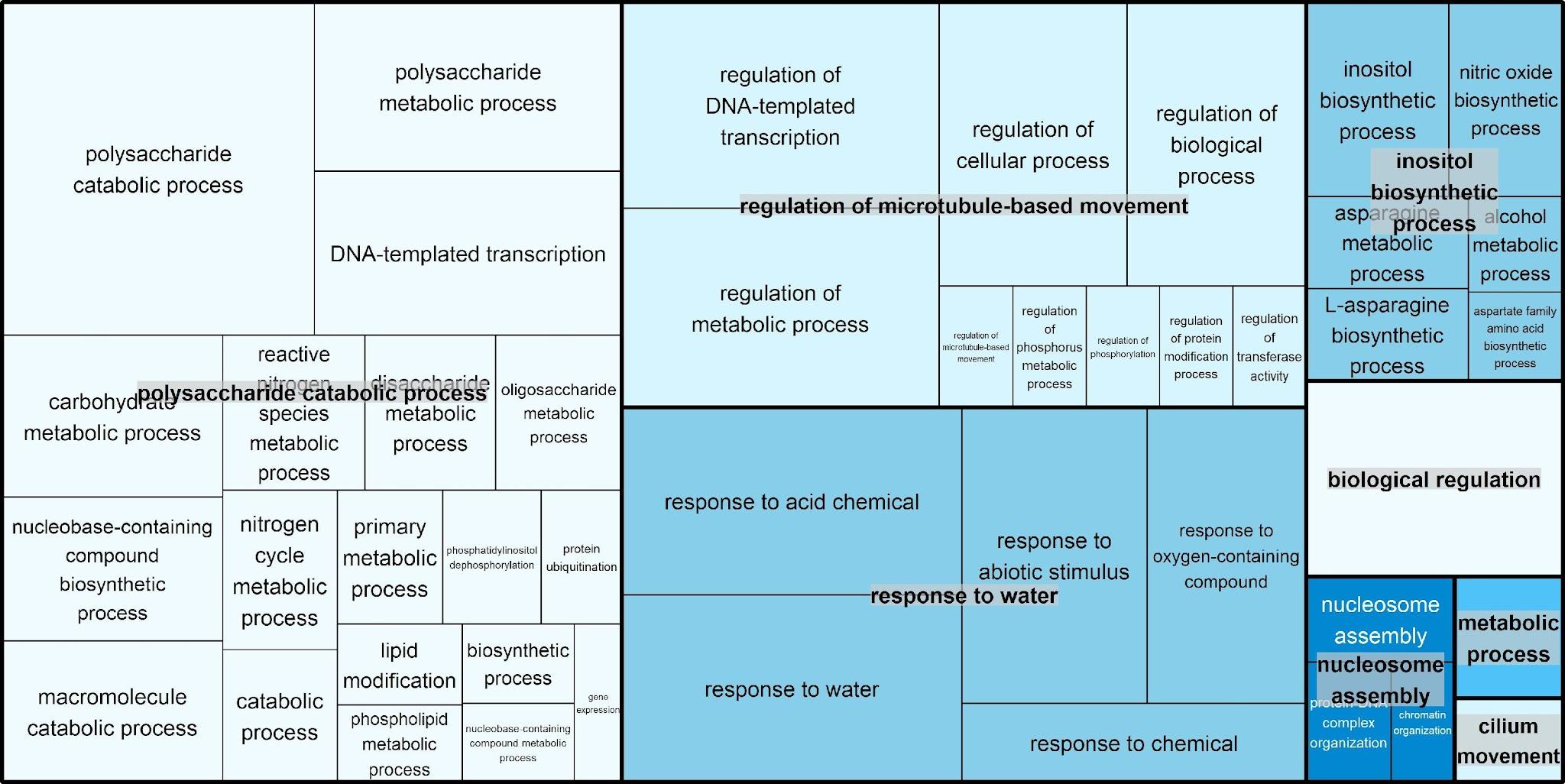

**i)**

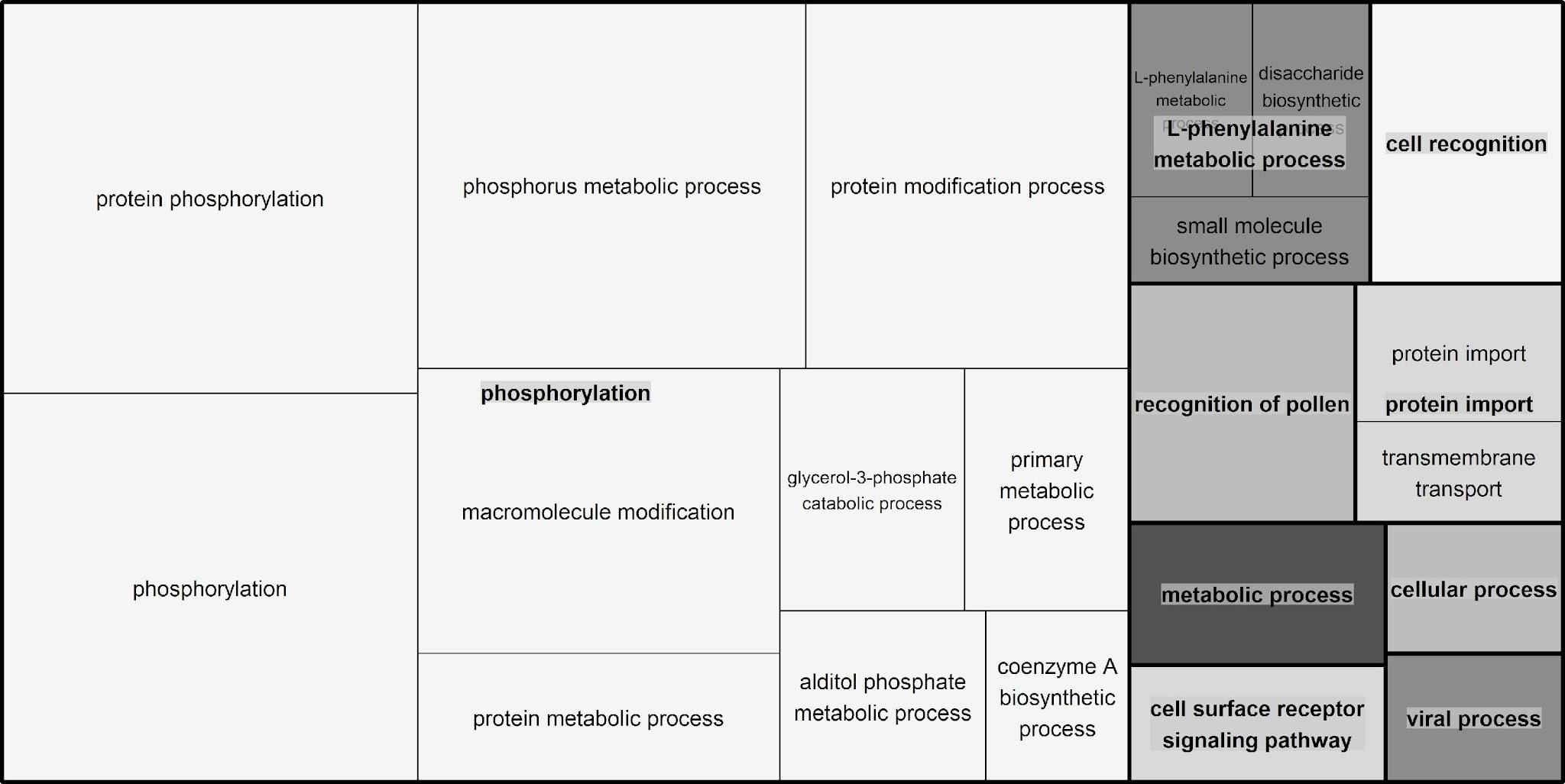

**Fig. S11. REVIGO TreeMaps summarizing Gene Ontology functional enrichment of co-expression modules showing overrepresentation of differentially expressed genes exclusive to smut-susceptible sugarcane genotypes.** Panels correspond to the following modules: (a) salmon, (b) steelblue, (c) black, (d) blue, (e) cyan, (f) midnightblue, (g) royalblue, (h) lightcyan, (i) darkolivegreen, and (j) violet. Rectangle size is proportional to the associated enrichment value.

**a)**

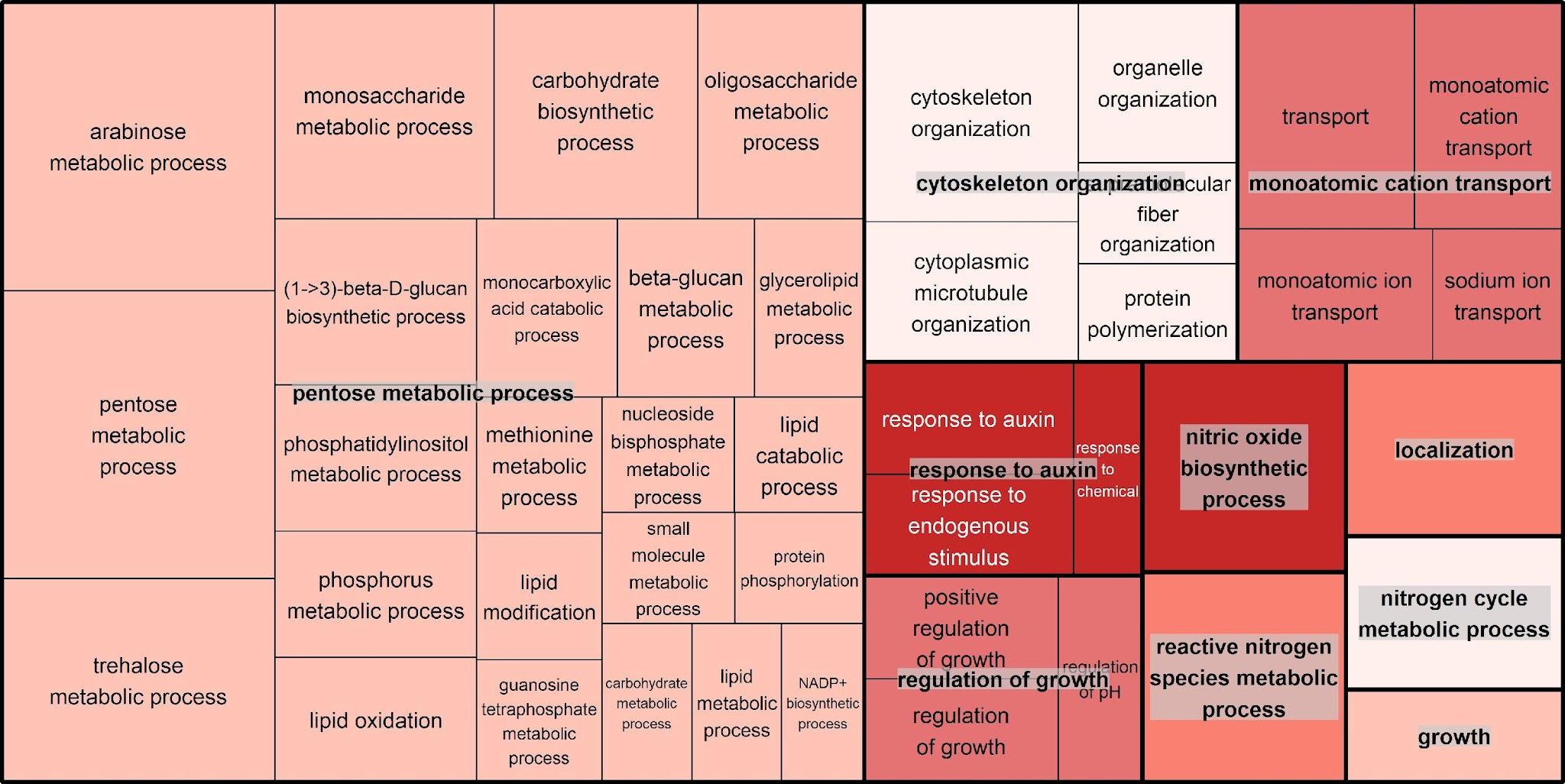

**b)**

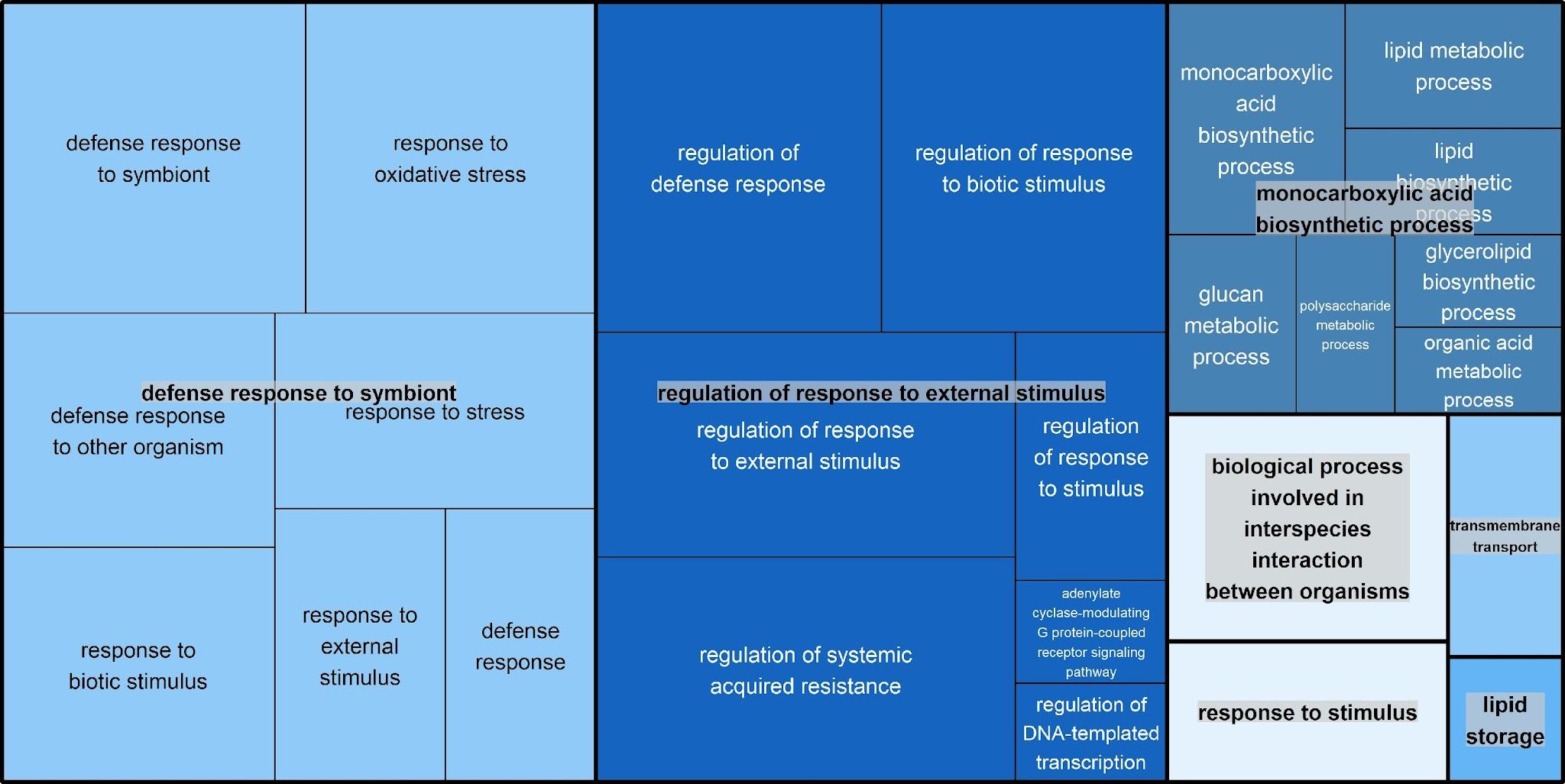

**c)**

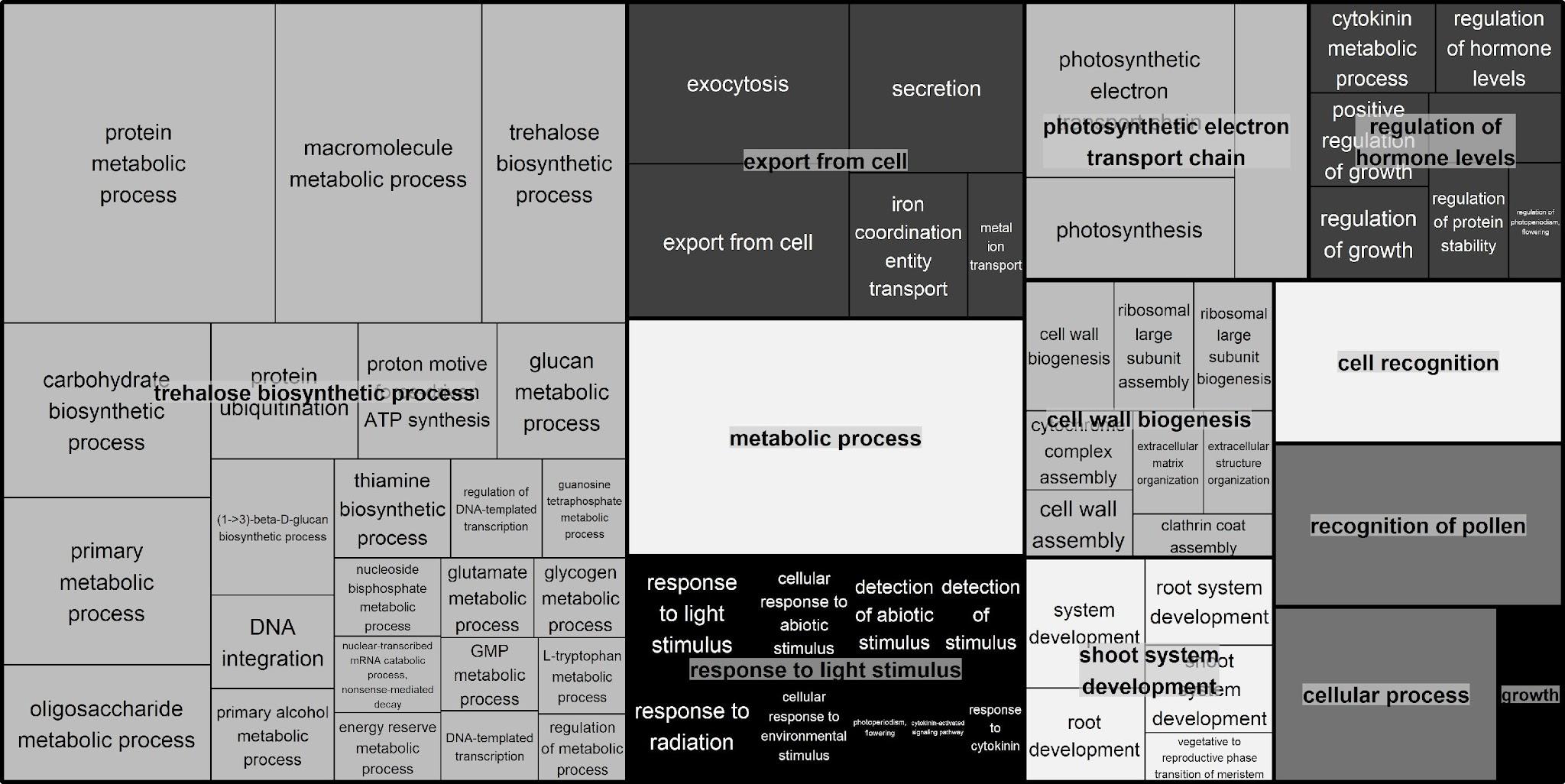

**d)**

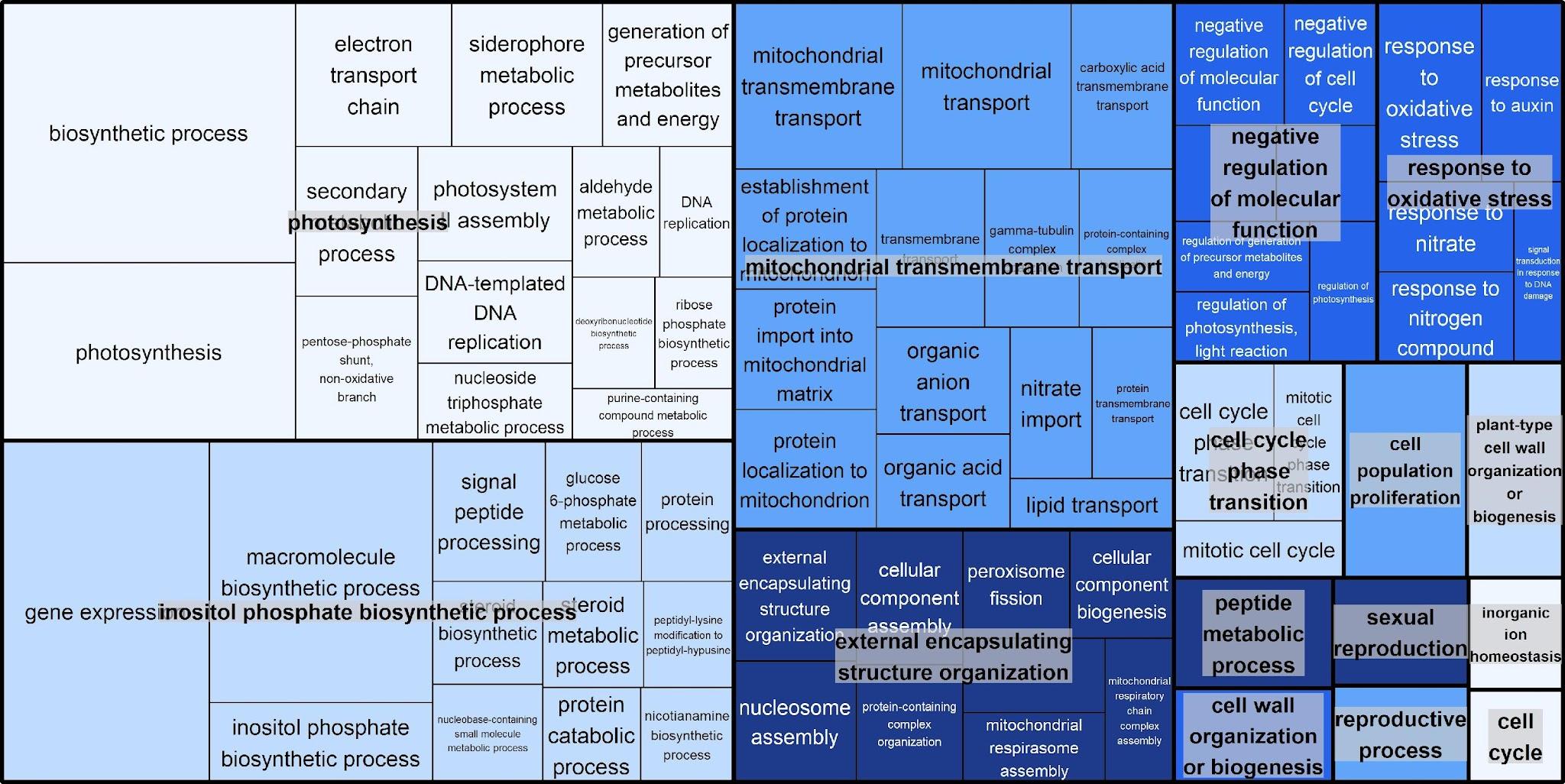

**e)**

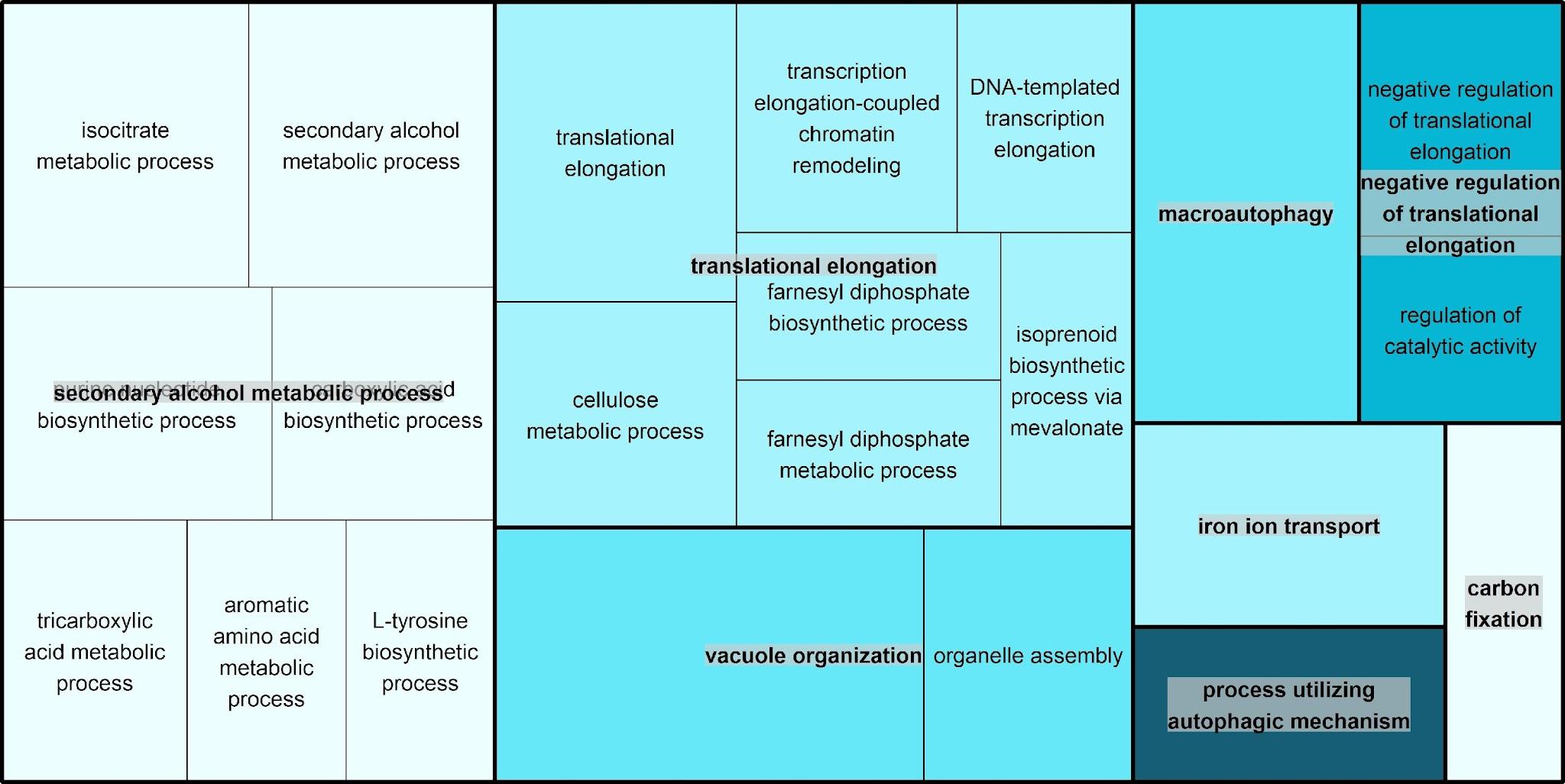

**f)**

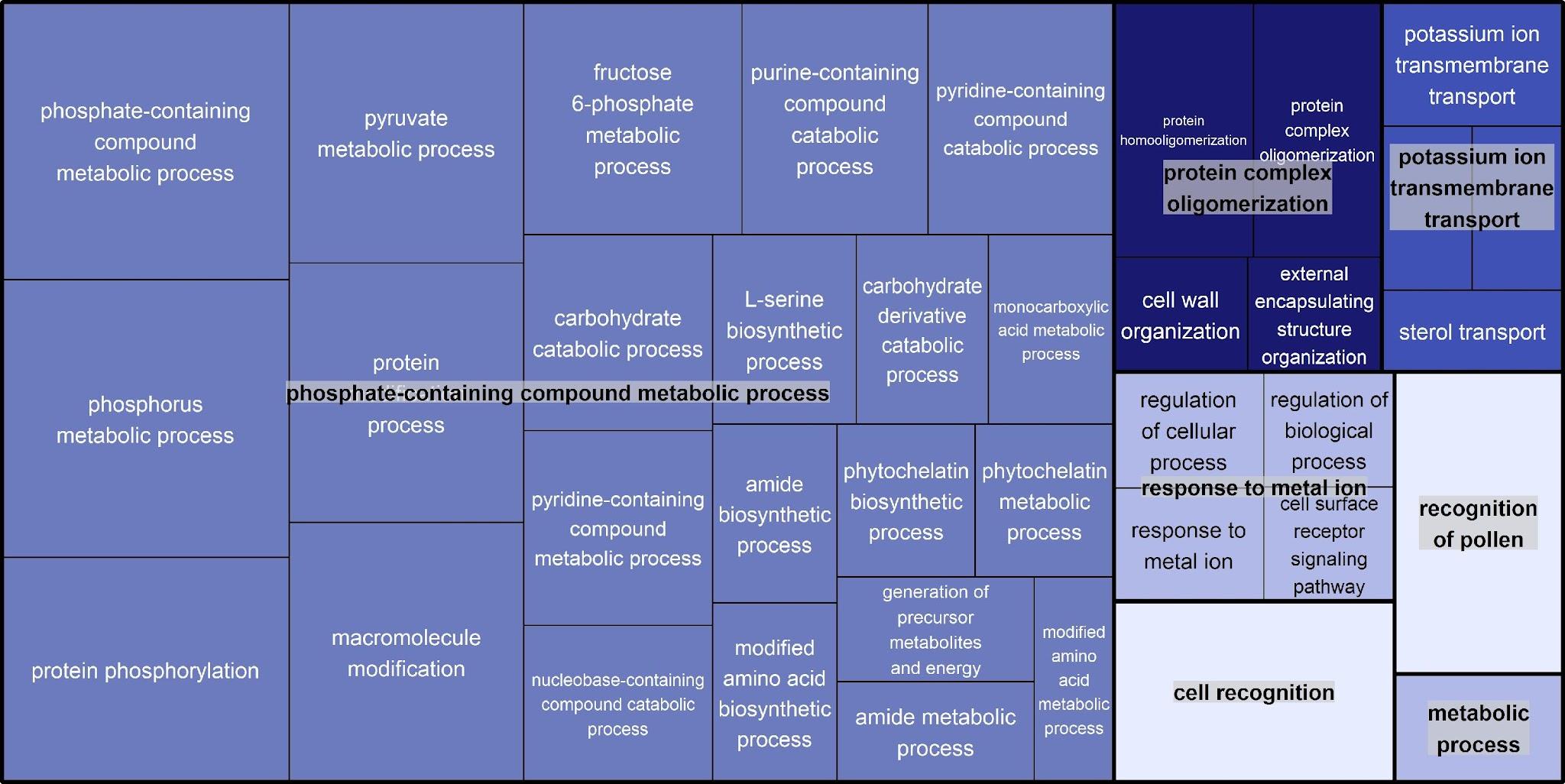

**g)**

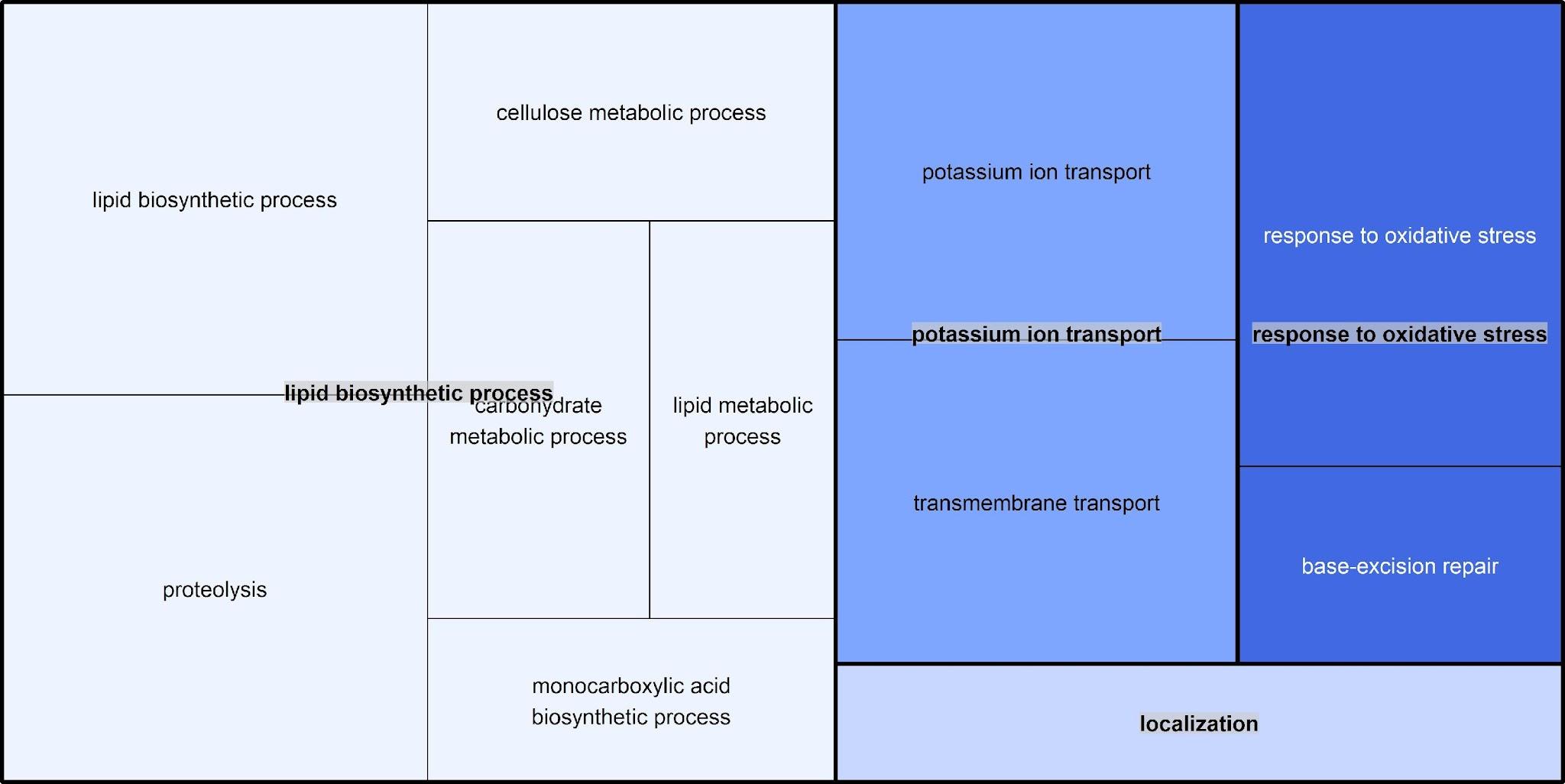

**h)**

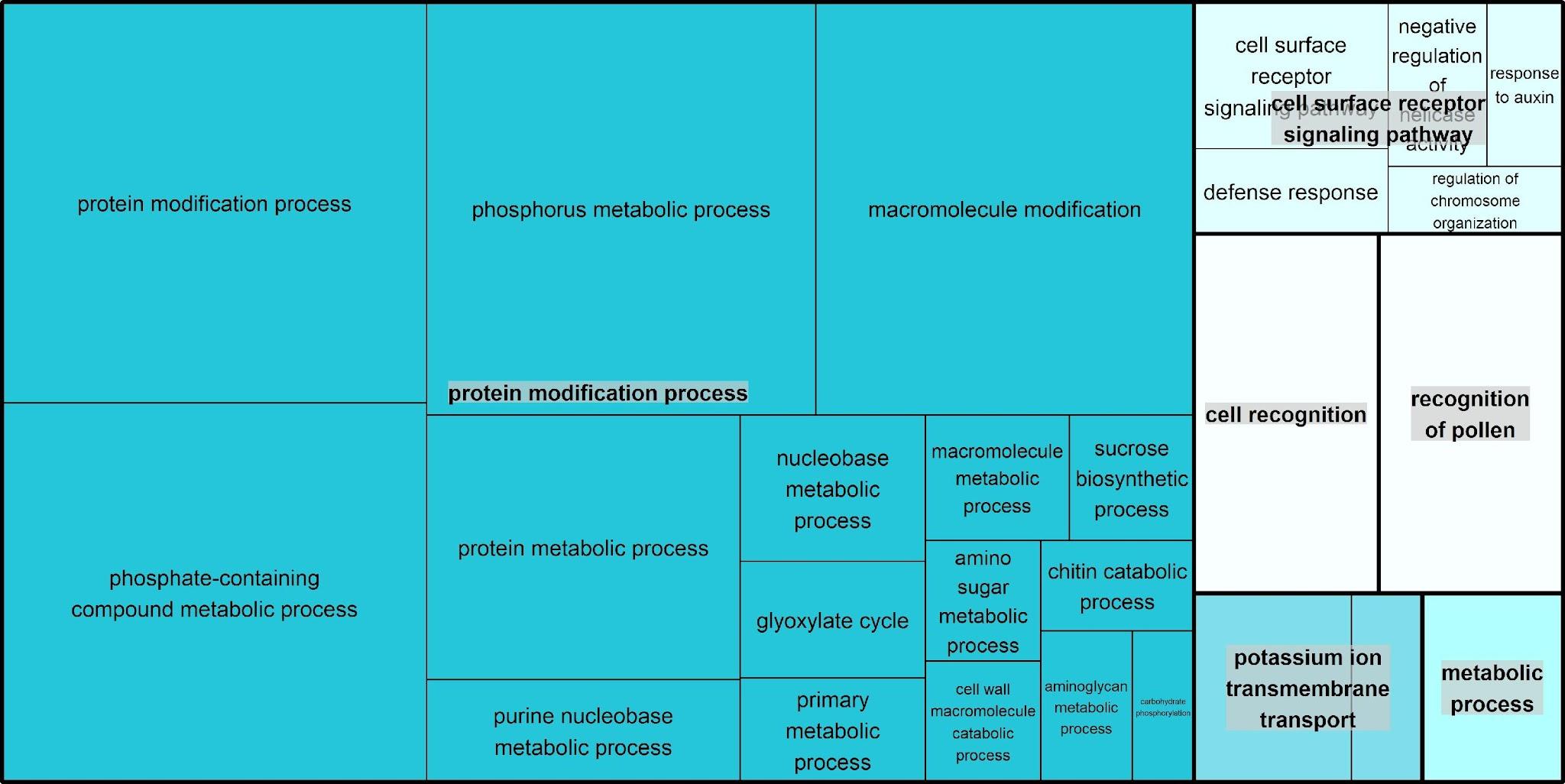

**i)**

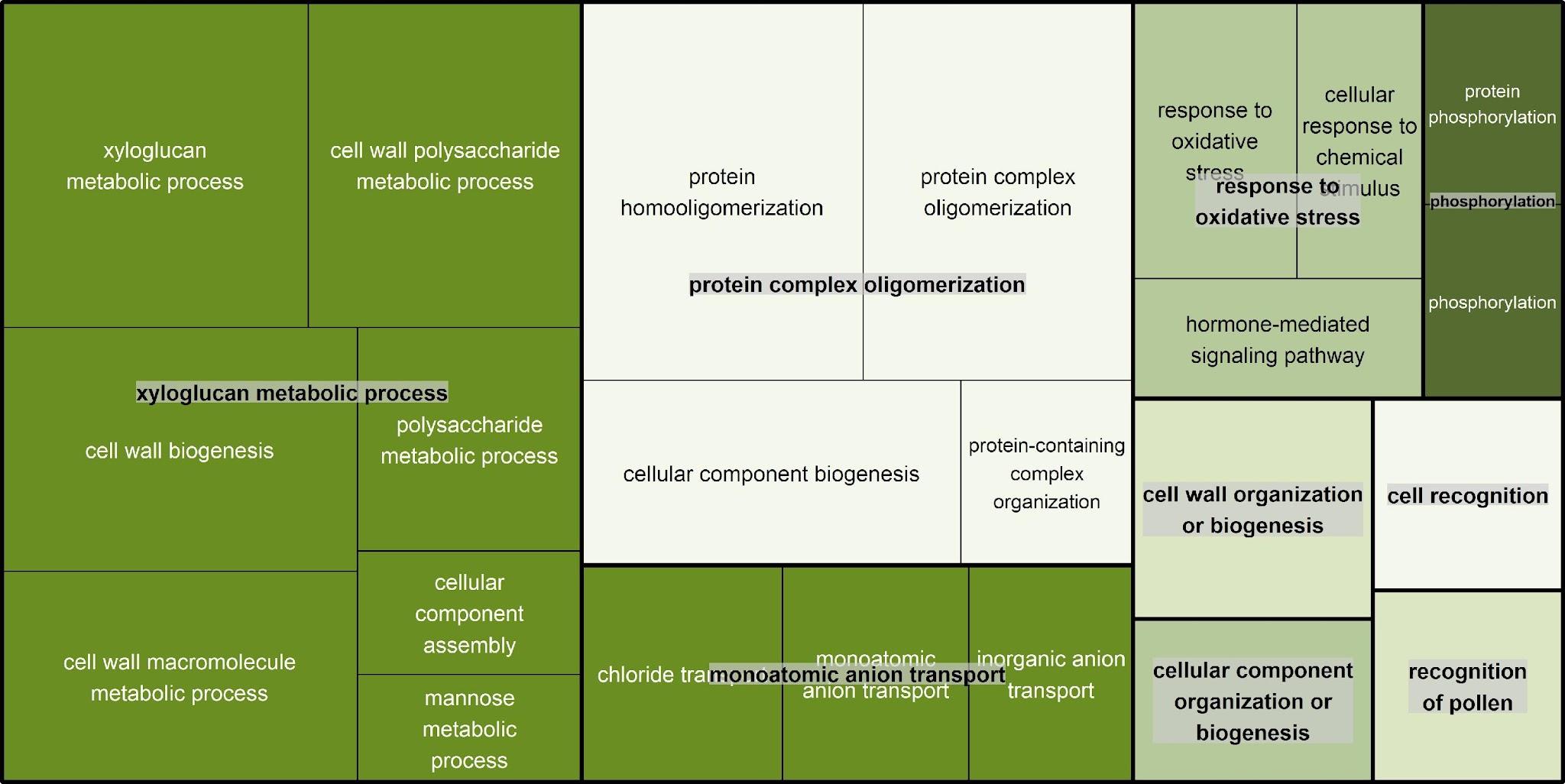

**j)**

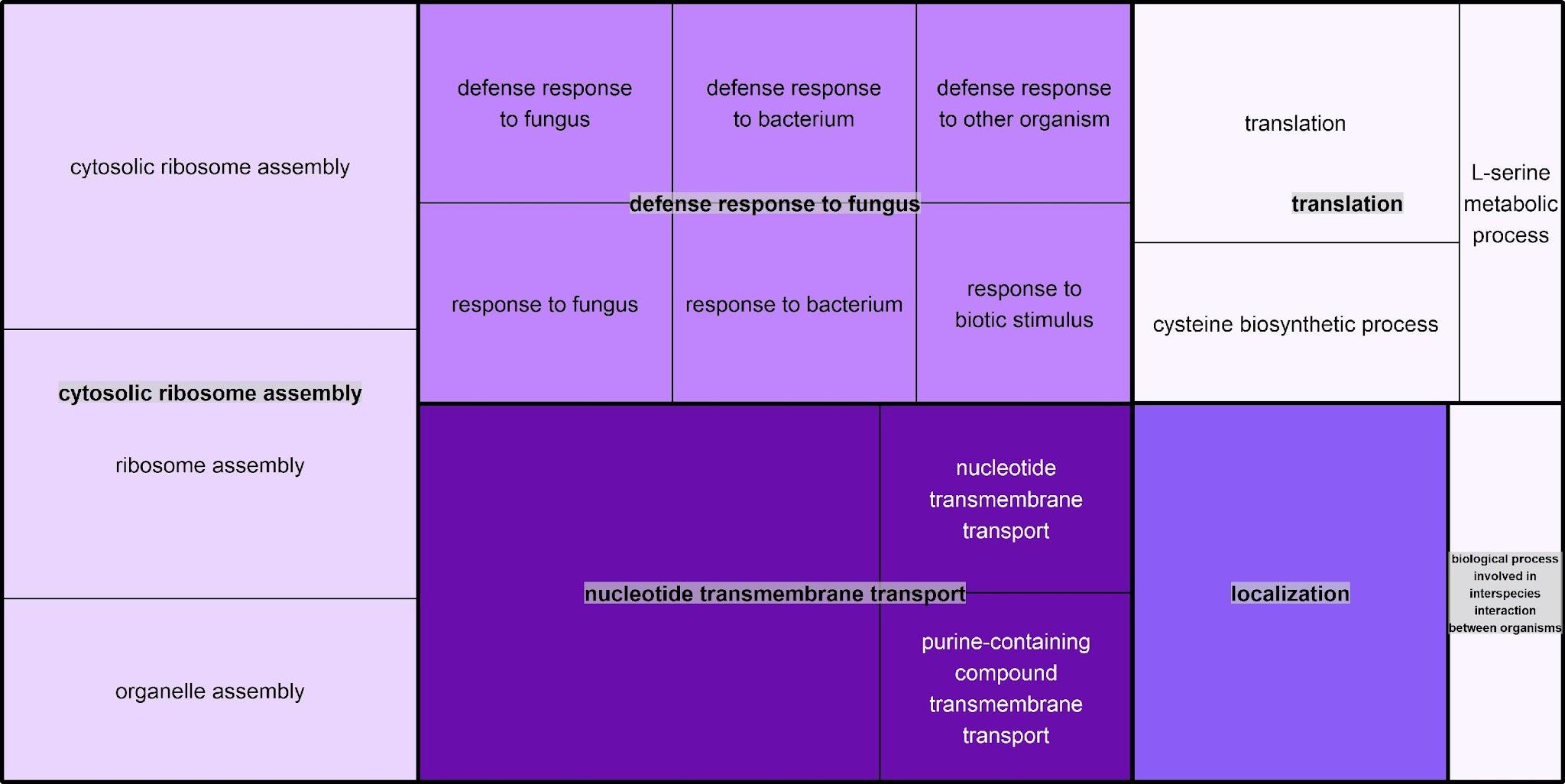
